# Network-aware reaction pattern recognition reveals regulatory signatures of mitochondrial dysfunction

**DOI:** 10.1101/2020.06.25.171850

**Authors:** Jordan A. Berg, Youjia Zhou, Yeyun Ouyang, T. Cameron Waller, Ahmad A. Cluntun, Megan E. Conway, Sara M. Nowinski, Tyler Van Ry, Ian George, James E. Cox, Bei Wang, Jared Rutter

## Abstract

Metabolism forms a complex, interdependent network, and perturbations can have indirect effects that are pervasive. Identifying these patterns and their consequences is difficult, particularly when the effects occur across canonical pathways, and these difficulties have long acted as a bottleneck in metabolic data analysis. This challenge is compounded by technical limitations in metabolomics approaches that garner incomplete datasets. Current network-based tools generally utilize pathway-level analysis lacking the granular resolution required to provide context into the effects of all perturbations, regardless of magnitude, across the metabolic network. To address these shortcomings, we introduce algorithms that allow for the real-time extraction of regulatory patterns and trends from user data. To minimize the impact of missing measurements within the metabolic network, we introduce methods that enable complex pattern recognition across multiple reactions. These tools are available interactively within the user-friendly Metaboverse app (https://github.com/Metaboverse) to facilitate exploration and hypothesis generation. We demonstrate that expected signatures are accurately captured by Metaboverse. Using public lung adenocarcinoma data, we identify a previously undescribed multi-dimensional signature that correlated with survival outcomes in lung adenocarcinoma patients. Using a model of respiratory deficiency, we identify relevant and previously unreported regulatory patterns that suggest an important compensatory role for citrate during mitochondrial dysfunction. This body of work thus demonstrates that Metaboverse can identify and decipher complex signals from data that have been otherwise difficult to identify with previous approaches.

---

Human metabolism has been elementally understood and appreciated since the time of Aristotle and the publication of *On the Parts of Animals* [1]. Over the past several centuries, metabolism has been further vigorously dissected to create a systematic map of metabolic reactions and their substrates, products, and modifiers. More recently, the study of metabolism has been aided immensely by the advent of high-throughput transcriptomics, proteomics, and metabolomics. Large consortia projects, such as the Kyoto Encyclopedia of Genes and Genomes (KEGG) [2,3], the Human Metabolome Database (HMDB) [4], and the Reactome Pathway Database [5–7], have also provided an invaluable and more holistic network perspective. Together, these resources have the potential to enhance our understanding of metabolism through enabling a more systematic query of datasets and databases. However, the complex and context-dependent nature of metabolism and the limited available tool set for metabolic pattern recognition and investigation from large datasets still confounds a more complete study of metabolism.

To circumvent the challenges related to metabolic complexity, it is common to adopt reductionist experimental approaches to tease apart the characteristics and mechanics of these processes and determine how they fit into the larger picture of biology and disease. Such strategies are vital to advancing our biological understanding but can also miss essential and multi-dimensional properties of metabolism. For example, in differential gene expression analysis, researchers rely on thresholds of magnitude and statistical significance to prioritize a short list of genes for further study. However, biological perturbations lead to complex, cooperative effects, many of which may seem negligible in isolation. Because metabolism is an interdependent system and distal components can have coordinated and rippling effects across the network, reductionist strategies can inadvertently limit the scope of a metabolism study [8, 9]. An additional challenge arises in metabolomics data analysis as it is common to only identify 100-200 metabolites for a given experiment [10]. This limitation leads to missing measurements across a given metabolic pathway, and can confound downstream data interpretation.

Several computational tools have risen to prominence over the past decade to try to resolve these issues in data analysis and interpretation; however, they cannot rapidly extract a comprehensive list of regulatory patterns within user data [5–7, 11–20] (see Suppl. Text 1 for further discussion of these tools). These shortcomings are particularly pronounced when experimental data have sparse coverage across the metabolic network. Without a more holistic integration of data on the metabolic network and the necessary capabilities to distil patterns across the metabolic network, many relevant mechanistic and regulatory metabolic patterns will be missed.

To address these limitations in metabolic data analysis, contextualization, and interpretation, we created Metaboverse, a cross-platform, interactive desktop application to aid users in exploring and interpreting their data’s metabolic story and generating new data-driven hypotheses. Metaboverse curates a reaction network database based on a Reactome knowledgebase [5–7], BiGG [21], or BioModels [22, 23] network. The user’s transcriptomics, proteomics, and/or metabolomics data can then be integrated into the reaction network in the form of log_2_(fold change) and statistical values for each measurement (as table rows) for each sample comparison (as table columns) (Fig. 1). Additional methods allow for the interpolation of protein complex measurements or protein measurements from upstream components. Once data are integrated onto the network, user-friendly interactive tools are available to visualize and explore reactions and their components individually, by canonical pathway definitions, or by nearest reaction neighborhood networks for a selected reaction and all its connected reactions, regardless of metabolic pathway participation. These flexible, visual approaches allow for less biased analysis by not constraining the user’s analysis to familiar pathways. This approach thus enables the exploration of patterns that may not be obvious when viewing canonical pathway depictions (Supp. Fig. 1; Supp. Fig. 2). Metaboverse and its documentation and supporting tutorials are available at https://github.com/Metaboverse and https://metaboverse.readthedocs.io.

**Fig 1.**
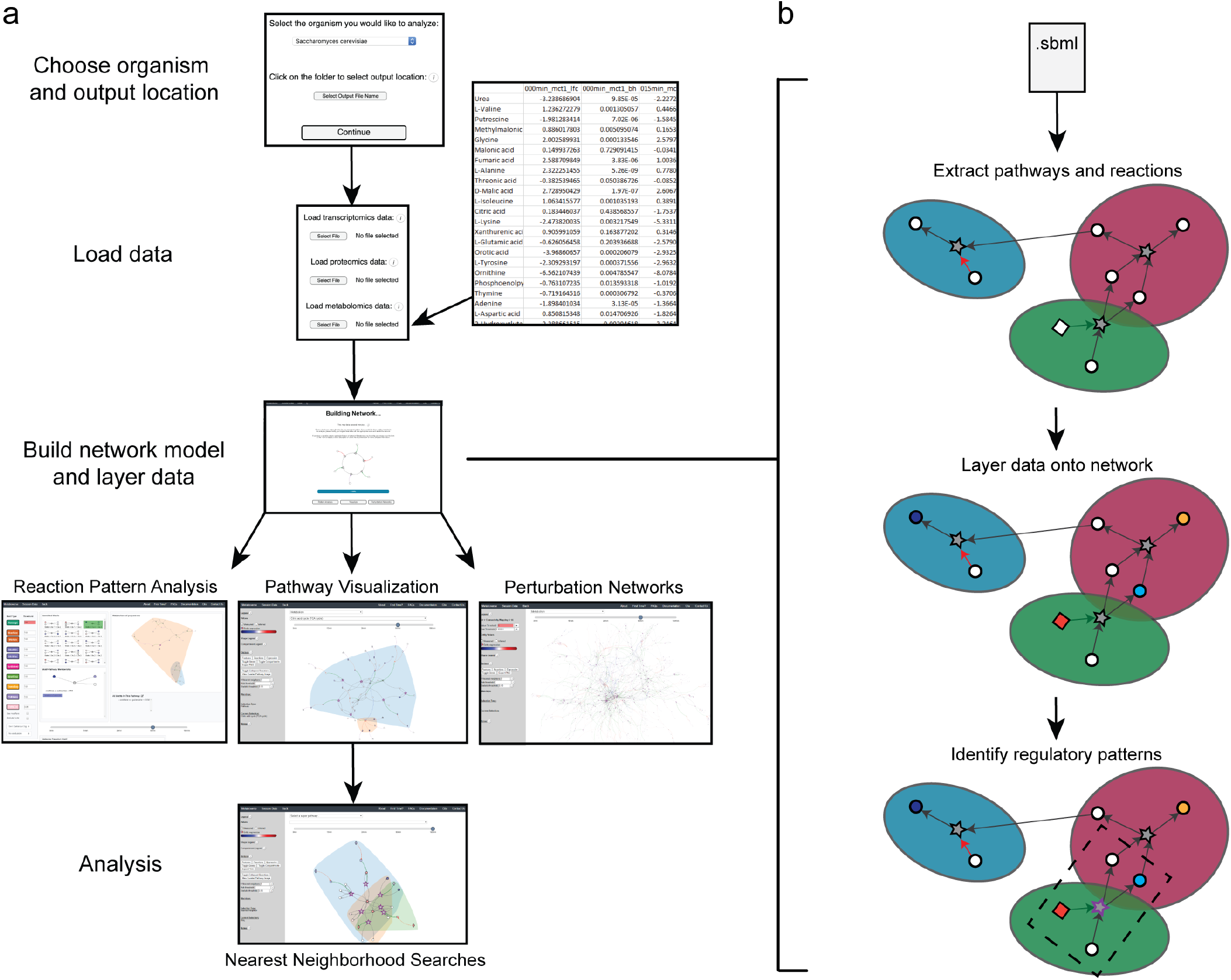
Metaboverse provides a simple, dynamic user interface for processing and exploring multi-omics datasets. **a**. The user provides the name of the organism of interest from a drop-down menu along with an output location. The user then has the option to provide transcriptomics, proteomics, and/or metabolomics datasets. These datasets can be single- or multi-condition or time-course experiments. Data is formatted as follows: row names are the measured entity names or IDs, the first column is a log_2_(fold change) or other measurement value, and the second column is a statistical measurement. For time-course and multi-condition datasets, this pattern is repeated for each subsequent sample as part of the same data table. During this step, the user can also provide sample labels and other modifiers to customize the curation and display of the data on the curated reaction network. Metaboverse will then build the model. Once the model is complete, the user will be able to visualize the patterns identified within reactions, explore pathway-specific or general perturbation networks, and perform general pathway and nearest reaction neighborhood exploration of the data. **b**. Overview of back-end metabolic network curation and data layering.

Tools that utilize user data and the metabolic network are generally limited to pathway-level analysis, or require the user to manually search for interesting patterns after projecting their data onto the network image [5–7, 11–20] (see Suppl. Text 1). Previous work by Checkik, et al. introduced the concept of an “activity motif”, where a network pattern was based on the expression characteristics of sequential components in a signaling cascade [24]. We previously identified multi-dimensional reaction-based patterns using a more manual approach [25], but based on similar principles. However, manually identifying a reaction-based pattern using multiple data measurements is time-consuming. Other network-based analysis tools often limit themselves to searching for structural network patterns and have been used with success predominantly for the analysis of protein-protein interaction networks. While these methods are useful for interaction networks, they are less tractable for the analysis of the metabolic network, where the graph structure is set and consistent, and the behavior of the network’s entities is where the user’s interest likely lies.

Critically, Metaboverse introduces, to our knowledge, the first implementations of algorithms that enable the rapid and automated discovery of complex regulatory patterns across all annotated reactions within that organism’s reaction network and across multiple sequential reactions. Metaboverse uses the metabolic reaction network to rapidly analyze each reaction on the fly for a variety of possible patterns based on the available transcript, protein, and metabolite measurements provided by the user (Fig. 2a). As an example, the Average reaction pattern search compares the averaged measurements of the reaction inputs and the averaged measurements of the reaction outputs to determine if there is some net change across the reaction. Metaboverse can also integrate information about a reaction’s modifying components, such as catalysts and inhibitors, to identify more complex patterns. Metaboverse then returns an interactive list of reactions that pass the predetermined fold change or statistical threshold for the user to explore (Supp. Fig. 3). A complete and current list and description of all available reaction pattern modules can be found in the documentation at https://metaboverse.readthedocs.io.

**Fig 2.**
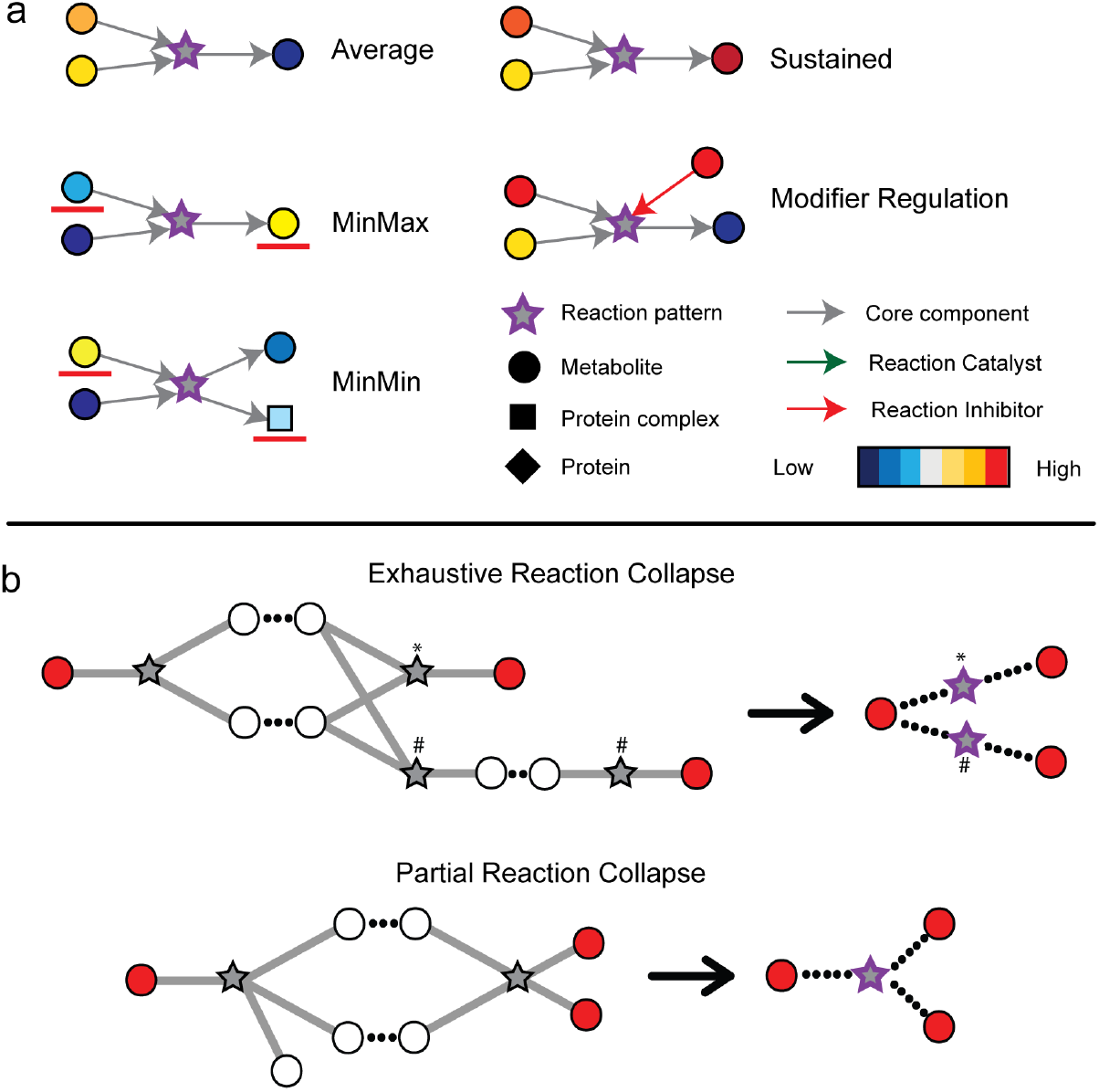
Overview of reaction pattern construction and reaction collapsing. **a**. Examples of a selection of reaction patterns available in Metaboverse. Reactions are depicted as stars, metabolites as circles, protein complexes as squares, and proteins as diamonds. Core interactions (inputs, outputs) are depicted as grey arrows, reaction catalysts as green arrows, and reaction inhibitors as red arrows. Component measurements are depicted in a blue-to-red color map, where lower values are more blue and higher values are more red. **b**. Example sub-networks where a reaction collapse would occur. Measured components are depicted as red circles, unmeasured components as white circles, and reactions as stars. Core interactions (inputs, outputs) are depicted as grey lines and identical components that would form the bridge between two reactions are depicted as dashed black lines between circles. A collapsed reaction is depicted as a star with a dashed border and its new connections between measured components are dashed black lines between a measured component and a reaction node. Collapsed reactions representing a particular reaction sequence are marked by an asterisk (*) or a number sign (#).

Missing data points, particularly in metabolomics experiments, are frequent and make the analysis of pathways and identification of regulatory patterns in the network challenging [10]. Thousands of metabolites are known to participate in human metabolism, yet the current state of metabolomics technologies is such that only a limited number of the metabolites are quantified. This limitation leads to gaps in the measured metabolic network and can confound pattern recognition across reactions. Therefore, we developed a reaction compression algorithm that collapses up to three connected reactions with intermediate missing data points if they can be bridged with measured data on the distal ends of the reaction series (Fig. 2b). Similar concepts have been used successfully in metabolomic analysis to define amino acid-related metabolites [26]. To our knowledge, Metaboverse provides the first computational and automated implementation for creating summarized reaction representations.

To demonstrate the ability of Metaboverse to identify known and new metabolic regulatory patterns, we first turned to public steady-state metabolomics data from early stage human lung adenocarcinomas [27]. In the original study of this dataset, the authors asked which metabolites could be used as diagnostic markers to identify early stage adenocarcinomas. It is important to identify reliable markers for this pervasive and deadly disease to provide viable and timely treatment options to patients. However, at the time of publication of the original study, available screening methods were prone to false positives. We asked if Metaboverse could capture the metabolic perturbations identified in this study, and identify other, more complex metabolic patterns.

Consistent with the original study [27] and our recent manual re-analysis of the data [25], steady-state abundances of nucleotide metabolism components were broadly upregulated in adenocarcinomas. Reaction pattern recognition by Metaboverse within and across canonical pathway representations, and using compressed reaction representations, reliably captured perturbations to nucleotide metabolism as the top hits (Supp. Fig. 4). Additionally, Metaboverse identified a reaction pattern that related to the perturbation of xanthine, a metabolite highlighted in the original study [27] (Supp. Fig. 5a). Interestingly, of the measured and statistically significant metabolite fold changes involved in the TCA cycle, we observed a reaction pattern driven by a reduced relative abundance of citrate and an increased relative abundance of malate (Supp. Fig. 5b). The changes in the concentrations of these two metabolites were both identified in the original study, but no discussion of their possible roles in this disease cohort, nor the fact that they are connected in the metabolic network, are found in the original study [27]. However, these metabolites are both components of a pathway that is integral to cancer metabolism. Their perturbations and interactions may provide valuable insights into the metabolic nature of lung adenocarcinomas. Metaboverse focuses the user’s attention to these metabolites and specifically their relationships to one another.

As we previously emphasized [25], by comparing the measured substrate and products of a reaction, we can identify multidimensional regulatory patterns that provide further insights into metabolic behavior. Strikingly, the top-ranking reaction pattern identified by Metaboverse involved the enzymatic conversion of dc-adenosyl methionine and spermidine to form spermine and 5′-methylthioadenosine by Spermine Synthase (SMS) (Fig. 3a). This reaction is connected to both polyamine synthesis pathway and the urea cycle. The second top-ranking reaction pattern centered around glycerate kinase (Fig. 3b). This reaction pattern, which consisted of a modest increase of measured glyceric acid coupled with the decrease of 3-phosphoglyceric acid, was identified manually in our previous reanalysis of the data [25], and could indicate a change in activity of glycerate kinase (GLYCTK). These changes across the GLYCTK reaction could be consequential as the reaction has connections to serine metabolism, a pathway that contributes to generating an ideal biosynthesis environment for tumorigenesis [28]. These connections were missed in the original study as a reaction-level analysis was not performed and these metabolites’ behavior in relation to each other was not emphasized. However, the significance of these reaction patterns was obvious with Metaboverse’s reaction pattern analysis module and these patterns were quickly and automatically highlighted to the user for further investigation.

**Fig 3.**
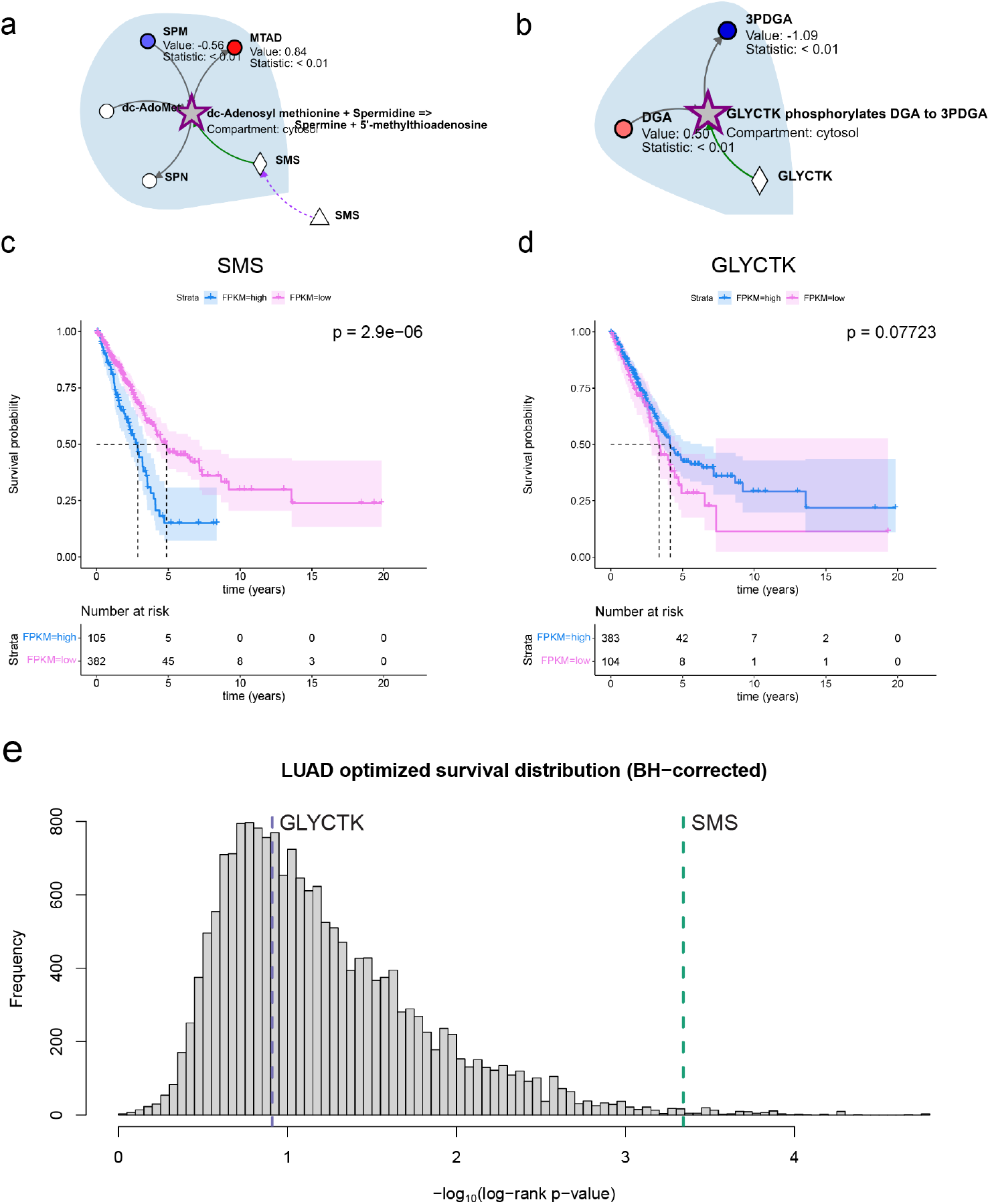
Metaboverse identifies novel putative regulatory signatures in early stage lung adenocarcinoma steady-state metabolomics data. **a**. Reaction between spermidine and 5′-methylthioadenosine identified by Metaboverse’s pattern recognition analysis module as the first-highest ranking `Average` reaction pattern. **b**. Reaction between glyceric acid and 3-Phosphoglyceric acid identified by Metaboverse’s pattern recognition analysis module as the second-highest ranking `Average` reaction pattern. Kaplan-Meier plots for the optimal expression cut-offs calculated for **c**. SMS (Spermine Synthase, FPKM cut-off: 49.5413; high: 105 tumors, low: 382 tumors) and **d**. GLYCTK (Glycerate Kinase, FPKM cut-off: 0.913; high: 104 tumors, low: 383 tumors). Shading indicates 95% confidence intervals for each expression group. Dashed lines indicate median survival times for each expression group. Risk tables are displayed below each Kaplan-Meier plot, and include the number of individuals in each risk category at time = 0 years. **e**. Distribution of Benjamini-Hochberg-corrected log-rank p-values for the Cox proportional hazards regression for each gene in the TCGA lung adenocarcinoma RNA-sequencing cohort. The rankings for SMS and GLYCTK are denoted by the dashed and labeled vertical lines. Metabolomics values are shown as node shading, where an increasingly blue shade indicates downregulation, and an increasingly red shade indicates upregulation. Measured log_2_(fold change) and statistical values for each entity are displayed below the node name. A gray node indicates a reaction. A bold gray node with a purple border indicates a motif at this reaction. Circles indicate metabolites and diamonds indicate proteins. Gray edges indicate core relationships between reaction inputs and outputs. Green edges indicate a catalyst. Background shading demarcates different cellular compartmentalization.

We initially explored these patterns using tools available within The Protein Expression Atlas [29–31] (v19.3) and discovered striking correlations between the gene expression of the enzymes mentioned above and patient survival outcomes in the lung adenocarcinoma (LUAD) cohort from The Cancer Genome Atlas (TCGA). We therefore performed Cox (Proportional Hazards) regression analysis for each gene in the TCGA cohort using an optimized expression cut-off for each gene [32–34]. SMS expression showed a remarkable correlation between high- and low-expression and patient outcomes (optimized FPKM cut-off: 49.5413; Fig. 3c,e; Supp. Fig. 6). Notably, the log-rank p-value for this relationship between patient outcome and SMS gene expression ranked in the top 0.65% of all regressions (#118 / 18169 surveyed genes). The log-rank p-value for GLYCTK in early stage adenocarcinomas was poor, and is potentially explained by the lower expression levels of GLYCTK in this cohort, thus is it difficult to draw a clear connection between the metabolomics and gene expression data in this case (optimized FPKM cut-off: 0.913; Fig. 3d,e; Supp. Fig. 6).

One point worth emphasizing is the predicted directionality of each enzymatic reaction based on the metabolomics data and their corresponding survival prediction. For the reaction catalyzed by SMS, higher expression of SMS correlated with poorer prognosis. This correlation was also reflected in the observations made from the metabolomics data where adenocarcinoma tissue had decreased spermidine concentrations and increased 5′-methylthioadenosine concentrations compared to normal tissue. While caveats exist in how we interpret steady-state metabolomics data, we could reasonably infer that SMS is more active or abundant in early stage lung adenocarcinomas and is thus generating more reaction product. This metabolite flux could then feed polyamine synthesis and fuel tumorigenesis and/or proliferation. One recent study in colon adenocarcinomas has implicated a role for SMS in the silencing of *Bim*, which encodes a pro-apoptotic factor, thus promoting cell survival [35]. Similar patterns in SMS have also been identified in mouse xenografts of lung cancer [36]. There has been little literature-based evidence for a role of polyamine metabolism, much less SMS, in lung adenocarcinomas. While the involved metabolites were singly identified in the original study by their fold changes and a connection to polyamine synthesis is mentioned [27], no context as to their reaction-level roles and consequences in lung adenocarcinomas is provided. Metaboverse quickly provides this resolution and multidimensional context and places a renewed emphasis on the need for further research into SMS’s role in lung adenocarcinomas and tumor proliferation. Conversely, while statistical power was weak, lower GLYCTK expression correlated with poorer prognosis, which reflected the observations made in the steady-state metabolomics data where adenocarcinoma tissue had increased glyceric acid concentrations and decreased 3-phosphoglyceric acid concentrations compared to normal tissue, indicating this enzyme’s activity or abundance may be decreased in tumors compared to normal tissue. These data generate new hypotheses regarding the role of these enzymes in early stage lung adenocarcinomas, but further investigation should be performed to definitively assess their impact in this disease. Findings like these nevertheless provide important context for understanding the metabolic regulatory environment and its reprogramming within lung adenocarcinomas and may inform research directions and treatment strategies for this disease.

To further demonstrate the capabilities of Metaboverse, we analyzed a model of mitochondrial fatty acid synthesis (mtFAS) deficiency in *S. cerevisiae*. mtFAS is an evolutionarily conserved pathway that produces lipoic acid, a critical co-factor for several metabolic enzymes. Recent work has uncovered additional important biological roles for this pathway. For example, mtFAS coordinates fatty acid synthesis with the regulation of iron-sulfur (Fe-S) cluster biogenesis and the assembly of mitochondrial oxidative phosphorylation complexes [37–39]. The relatively recent discovery of patients with mutations in genes encoding key mtFAS enzymes further illustrates the importance of this pathway in human physiology [40]. The *S. cerevisiae* Mct1 protein, homologous to the Malonyl-CoA-Acyl Carrier Protein Transacylase (MCAT) in humans, transfers a malonyl group from malonyl-CoA to the mitochondrial acyl carrier protein (ACP). Deletion of the *MCT1* gene abolishes the activity of the mtFAS pathway in yeast [38]. We therefore used an *mct1*Δ mutant to explore the relationship between mtFAS pathway activity and the effects of its perturbation on downstream metabolic processes.

Previously, we generated proteomics data in *mct1*Δ yeast after the shift from fermentable to non-fermentable growth media [38]. We complemented these data with RNA sequencing at 0, 3, and 12 hours and steady-state metabolomics at 0, 0.25, 0.5, 1, 3, and 12 hours after this shift in growth media. By transitioning from a fermentable to a non-fermentable carbon source, yeast move from a glucose-repressed state to a state of increased respiratory potential. Upon layering these data on the *S. cerevisiae* metabolic network using Metaboverse and analyzing the network using the reaction pattern recognition module, we were able to explore acute and extended metabolic responses to mtFAS deficiency. Importantly, the top-ranked `ModReg` reaction patterns (by magnitude where both sides were statistically significant) at 12 hours predictably centered around Mct1 protein abundance, Coenzyme A species measurements, and other fatty acid synthesis-related reactions (Fig. 4a; Suppl. Fig. 7a-c,e-h).

**Fig 4.**
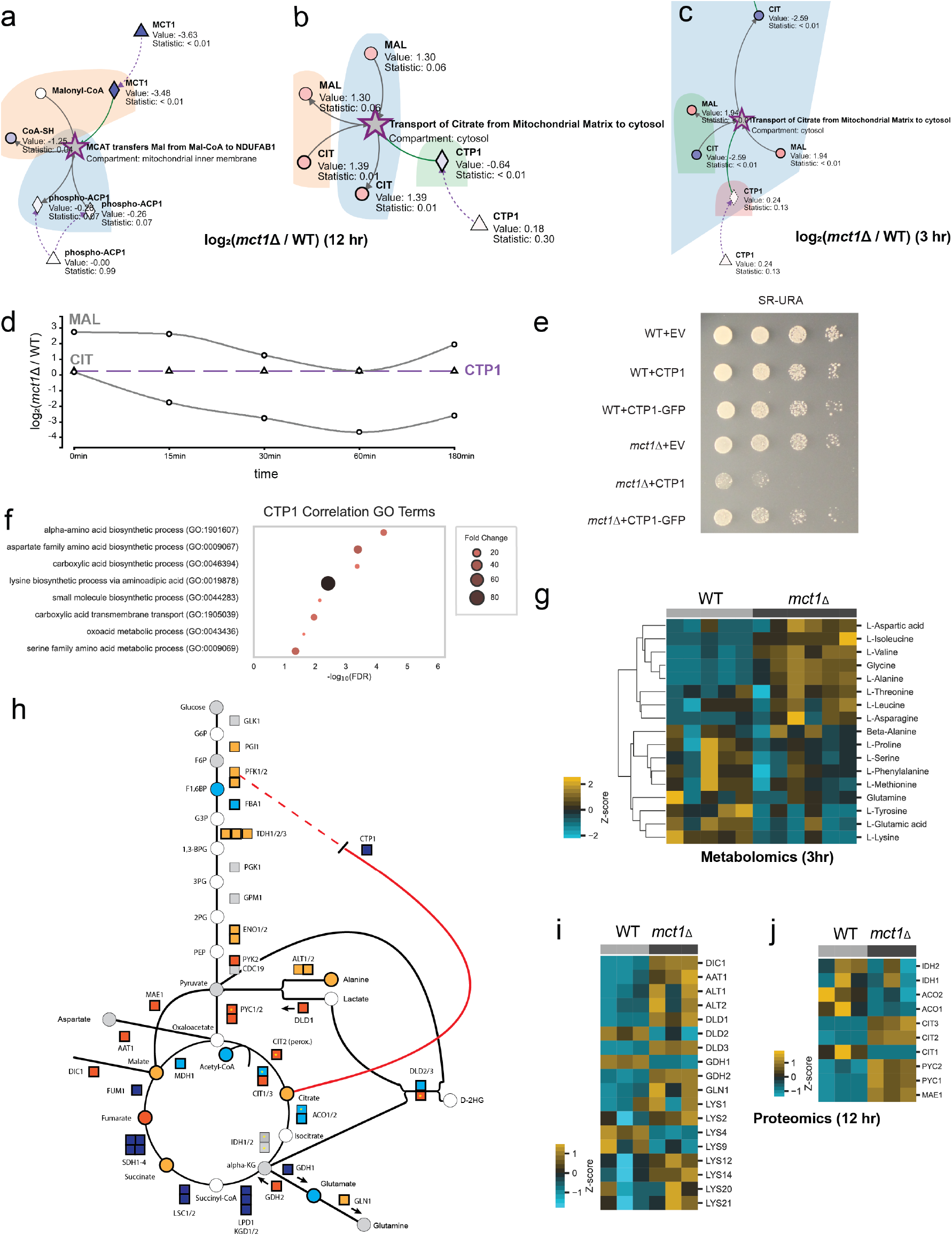
Metaboverse identifies novel compensatory mechanisms to mitochondrial dysfunction. **a**. “MCAT transfers Mal from Mal-CoA to NDU-FAB1” reaction pattern. **b**. “Transport of Citrate from Mitochondrial Matrix to cytosol” reaction pattern using steady-state metabolomics and proteomics data (12 hours). **c**. “Transport of Citrate from Mitochondrial Matrix to cytosol” reaction pattern using early (3 hours) metabolomics and RNA-sequencing data. **d**. Time-course behavior of citrate and isocitrate metabolites at earlier (0, 0.25, 0.5, 1, 3-hour) time-points. **e**. Spot growth assays for wild-type and *mct1*Δ. mutant strains with no expression (EV), overexpression of *CTP1*, and overexpression of *CTP1*-GFP construct on raffinose-supplemented growth medium. **f**. Bubble plot for the GO term enrichment results for genes identified in the SpQN-corrected co-expression analysis of *CTP1* in the refine.bio wild-type cohort (n = 1248). -log_10_(FDR) is plotted along the x-axis and fold change enrichment is plotted as bubble size and color intensity. **g**. Heatmap of amino acid metabolites for wild-type and *mc11*Δ. mutant strain proteomics at 3 hours post-raffinose carbon source shift. **h**. Graphical overview of yeast glycolysis pathway and other related reactions overlaid with summary annotations based on RNA-sequencing, proteomics, and metabolomics measurements during steady-state growth (12 hrs). Circles indicate metabolites and squares indicate proteins. RTG target proteins also contain a yellow asterisk within the square. Entities with no measurement are filled white, entities with no significant change are shaded grey. Increasingly yellow to orange shades indicate positive fold changes and increasingly blue shades indicate negative fold changes. A more complete legend for the shading criteria can be found in Supp. Fig. 13. **i**. Heatmap of amino-acid-regulated enzymes for wild-type and *mct1*Δ. mutant strain proteomics at 12 hours post-raffinose carbon source shift. **j**. Heatmap of anaplerotic enzymes for wild-type and *mct1*Δ. mutant strain proteomics at 12 hours post-raffinose carbon source shift. Measured values are shown as node shading in network plots, where an increasingly blue shade indicates downregulation, and an increasingly red shade indicates upregulation. Measured log_2_(fold change) and statistical values for each entity are displayed below the node name and represent the comparison of *mct1*Δ. vs. wild-type samples. RNA-sequencing comparisons contained n=4 in each group, proteomics comparisons contained n = 3 in each group, and metabolomics comparisons contained n = 6 in each each comparison group, except for the 3-hour wild-type group, which contained n = 5. A gray node indicates a reaction. A bold gray node with a purple border indicates a potential regulatory pattern at this reaction for the given data type time points. Circles indicate metabolites, diamonds indicate proteins, and triangles indicate gene components. Gray edges are core relationships between reaction inputs and outputs. Green edges indicate a catalyst. Dashed blue edges point from a metabolite component to the complex in which it is involved. Dashed orange edges point from a protein component to the complex in which it is involved. Dashed purple edges point from a gene component to its protein product. Protein complexes with dashed borders indicate that the values displayed on that node were inferred from the constituent protein, metabolite, and gene measurements. Background shading demarcates different cellular compartmentalization. Heatmap values were mean-centered at 0 (z-score). Hierarchical clustering was performed where indicated by the linkage lines using a simple agglomerative (bottom-up) hierarchical clustering method (or UPGMA (unweighted pair group method with arithmetic mean)).

We also observed respiratory signatures consistent with our previous studies [38, 41], including, for example, the identification of a statistically significant pattern in the electron transfer from ubiquinol to cytochrome C via Complex III of the electron transport chain (ETC) (Supp. Fig. 7g, 8a). At the protein level, the cytochrome C isoforms, Cyc1 and Cyc7, and Complex III components, were significantly decreased in abundance compared with wild-type cells. These components catalyze the transfer of electrons from ubiquinol to cytochrome C. We have observed this pattern in oxidative phosphorylation complexes during mtFAS perturbations by various other means [37, 38].

The second expected pattern of interest identified by reaction pattern analysis in Metaboverse was the general reduction of TCA cycle-related enzyme abundances and TCA cycle intermediate metabolite concentrations (Supp. Fig. 7c, 8b). We observed the upregulation of Dic1 protein abundance. Coincidentally, *DIC1* expression is essential for growth on non-fermentable media due to its role in shuttling phosphate across the mitochondrial inner membrane in exchange for malate or succinate [42]. These two metabolites are essential intermediates within the TCA cycle, and their depletion has negative consequences on mitochondrial respiration. When yeast are switched to a non-fermentable carbon source, especially when deficits in TCA cycle flux are present as in the mtFAS model, they appear to adapt by increasing Dic1 protein levels. This may in turn facilitate malate or succinate transport and aid in maintaining some level of TCA cycle flux and mitochondrial respiration. With this hypothesis in mind, it is interesting to see the general increase in whole-cell malate concentrations in the *mct1*Δ mutant compared to a wild-type strain.

One top-ranking, and frankly unexpected, reaction pattern identified using Metaboverse in the yeast multi-omics dataset was “Transport of Citrate from Mitochondrial Matrix to cytosol” (Fig. 4b; Suppl. Fig. 7d). This reaction pattern was also identified at the 3-hour time-point using metabolomics and RNA-sequencing data (Fig. 4c). We noticed that earlier metabolomics measurements showed citrate levels initially decreasing then increasing over the time-course (Fig. 4d). Ctp1, the protein catalyst governing this identified reaction pattern, is a tricarboxylate transporter that transfers citrate from the mitochondrial matrix to the cytosol [43].

Given that citrate is a key metabolite in the TCA cycle, we hypothesized that Ctp1 protein levels decrease in response to early citrate depletion to maintain citrate pools within the mitochondrial matrix, where it is perhaps most physiologically important for these cells to be able to adapt to the loss of *MCT1*. If *mct1*Δ cells downregulate Ctp1 as an adaptive mechanism, then forced overexpression of Ctp1 should lead to growth defects in this context. Indeed, we observed a specific sensitivity to Ctp1 overexpression in the *mct1*Δ background (Fig. 4e; Suppl. Fig. 14). This is despite elevated whole-cell concentrations of citrate in the *CTP1*-overexpression background compared to *mct1*Δ alone (Suppl. Fig. 9f). We also determined that the *CTP1* overexpression-GFP C-terminal fusion vector ablated this growth defect, suggesting the perhaps the GFP-fusion vector of Ctp1 was inactive and this growth phenotype was specific to functional Ctp1 localizing to the mitochondria and not spurious effects from protein overexpression.

We next performed co-expression analysis with the yeast transcriptomics compendia from refine.bio [44, 45]. We identified genes that correlated (r *>* 0.5) with *CTP1* expression in wild-type yeast across various experimental conditions. Enriched GO terms from this analysis revealed enrichment of aspartate, lysine, and other amino acid biosynthesis programs (Fig. 4f; Suppl. Fig. 10; Suppl. Fig. 11). Of note, aspartate can be converted into fumarate via the urea cycle and purine synthesis pathway, which might partially explain increased fumurate concentrations despite a non-functional electron transport chain. Additionally, lysine can be used as a substrate for the generation of acetyl-CoA, which plays a central role in the mtFAS pathway [38], and provides further evidence connecting *MCT1* back to mtFAS and *MCT1* deletion. These co-expression patterns were reflected in the yeast refine.bio compendium across all conditions and genetic backgrounds [44]. Metabolic rewiring of these and other amino acids additionally were apparent in the metabolomics data (Fig. 4g) and in the proteomics data (Fig. 4i). These data suggest that broad rewiring of biosynthesis pathways is occurring in the *mct1*Δ background, potentially partially explaining its ability to grow normally compared to wild-type (Fig. 4e). However, *CTP1* overexpression in this case appears to disrupt this biosynthetic rewiring, disabling the ability of *mct1*Δ cells to tune their growth with their genetic defect and related mtFAS and respiratory defects.

One well studied response to mitochondrial stress is the retrograde signaling pathway, which is known to act as a bridge for communication between the mitochondria and nucleus [46]. Previous evidence has suggested that one of the goals of the retrograde signaling pathway is to maintain α-ketoglutarate pools for biosynthetic processes. We also observed increased protein abundance levels in components of this anaplerotic pathway, namely Mae1, Pyc1, Pyc2, Cit2/Cit3, and Gdh2 (Fig. 4j). These enzymes sequentially catalyze the conversions of malate to pyruvate to oxaloacetate to citrate, respectively. *PYC1* and *CIT2* are also targets of the retrograde signaling pathway [46]. Work on the retrograde signaling response has also suggested a role for elevated Cit2 expression in maintaining the metabolite pools necessary for anabolic growth [46], which we also see in the *mct1*Δ background through general rewiring of biosynthetic components at earlier measured time-points (Fig. 4j; Suppl. Fig. 12). If metabolic compensation in *mct1*Δ cells requires increased citrate pools within mitochondria to maintain α-ketoglutarate pools for biosynthetic processes, *CTP1* downregulation could contribute to mitochondrial citrate homeostasis.

We observed several other patterns that would enable this compensatory biosynthetic reprogramming. The protein abundances of Dld3, another target of the RTG pathway, and Dld1, were significantly increased in the *mct1*Δ background (Fig. 4i). Dld3 catalyzes the conversion of 2-hydroxyglutarate and pyruvate to α-ketoglutarate and lactate, while Dld1 catalyzes the conversion of lactate to pyruvate. These reactions’ coordination could thus supplement α-ketoglutarate pools while additionally processing any available lactate in the environment to provide further substrate for the Dld3-catalyzed reaction (Fig. 4h,i; Supp. Fig. 13).

Another predicted role for *CTP1* in the *mct1*Δ background relates to the regulation of glycolysis. Cytosolic citrate is a potent inhibitor of phosphofructokinase, which is an important regulator of glycolysis [47, 48]. In the *mct1*Δ background, we see the broad increase in glycolytic enzyme abundances and can thus predict a degree of carbon flux through the TCA cycle. Ctp1 is also decreased in overall abundance, so we can infer that phosphofructokinase inhibition by citrate is limited. However, when *CTP1* is overexpressed in the *mct1*Δ background, citrate is likely saturating in the cytosol and inhibiting glycolysis, leading to the downstream disruption of carbon flux through the TCA cycle (Supp. Fig. 13). This point is reflected in the comparison of *CTP1*-overexpression in the *mct1*Δ background to *mct1*Δ alone, where we observed decreased abundances of the expected downstream central carbon and TCA metabolism intermediates, such as succinate (Suppl. Fig. 9j) and pyruvate (Suppl. Fig. 9d), in response to *CTP1* overexpression.

These data and the insights provided by Metaboverse implicate a novel putative role for changes in Ctp1 as part of a broader response to mtFAS deficiency and mitochondrial dysfunction to maintain carbon flux in a manner that allows for continued cellular growth. Specifically, these data demonstrate that *CTP1* overexpression limits cell growth in a model of mtFAS deficiency and that *CTP1* expression is generally coordinated with amino acid biosynthesis. *CTP1* could thus be acting as part of a broader metabolic program which serves to sustain growth in scenarios where mitochondrial homeostasis is disrupted. Ctp1 also likely aids in maintaining carbon flux through portions of the TCA cycle that contribute to biosynthesis, even though respiration is generally non-operational in this biological model due to the inability of these cells to assemble respiratory complexes. It has long been recognized that the signaling and stress responses to mtFAS defects and mitochondrial dysfunction are broad, and the current literature has likely only scratched the surface regarding these responses. These lines of evidence place Ctp1 and citrate homeostasis within this regulatory equation and further emphasize that complex and coordinated compensatory mechanisms respond to mitochondrial dysfunction to allow for continued cellular growth and survival.

Using Metaboverse, we identified canonical metabolic phenomena across various datasets, as well as novel putative regulatory patterns that could be further tested. We demonstrated that using public metabolomics data from lung adenocarcinomas, we could identify reaction patterns that corresponded with patient survival based on the gene expression of these reactions’ enzymes. We demonstrated that reaction pattern identification in a model of yeast mitochondrial dysfunction could identify relevant regulatory patterns and compensatory mechanisms. Critically, Metaboverse identifies biologically relevant patterns that remain hidden using existing analytical approaches. Metaboverse is provided as an easy-to-use visual tool so that users with no computational experience can easily explore their datasets, identify relevant reaction patterns, and generate new and compelling hypotheses. We expect Metaboverse to become a foundational analytical tool and augment the user’s experience when analyzing their own data or pre-existing datasets. Users will be able to frame their data in the context of the entire metabolic reaction network, discover new and interesting patterns, and design experiments with a more holistic mindset to learn how reaction patterns fit into the larger metabolic story of their model.

## Methods

For the current instructions and details of features, we refer the user to the Metaboverse documentation (https://metaboverse.readthedocs.io). Processed data from the yeast *mct1*Δ. analysis are available in distributed versions of Metaboverse (as `test_data.zip`) to act as a test dataset for users to familiarize themselves with the input data format requirements.

### Network curation

Biological networks are curated using the current version of the Reactome knowledgebase [5–7]. In particular, the pathway records and Ensembl- and UniProt-Reactome mapping tables are integrated into the network database for Metaboverse. Additionally, the ChEBI and The Human Metabolome databases are also referenced for metabolite synonym mapping to accept more flexible metabolite input nomenclature from the user [4, 49]. These data are used to generate a series of mapping dictionaries for entities to reactions and reactions to pathways for the curation of the total reaction network. Reaction annotations are additionally obtained from the Reactome knowledgebase [5–7]. As of the time of writing, users can also provide BiGG [21] and BioModels [22,23] networks; however, full support cannot always be guaranteed due to the more bespoke nature of some network models from these sources. The resulting curation file is output as a pickle-formatted `.mvdb` file.

**Listing 1.**
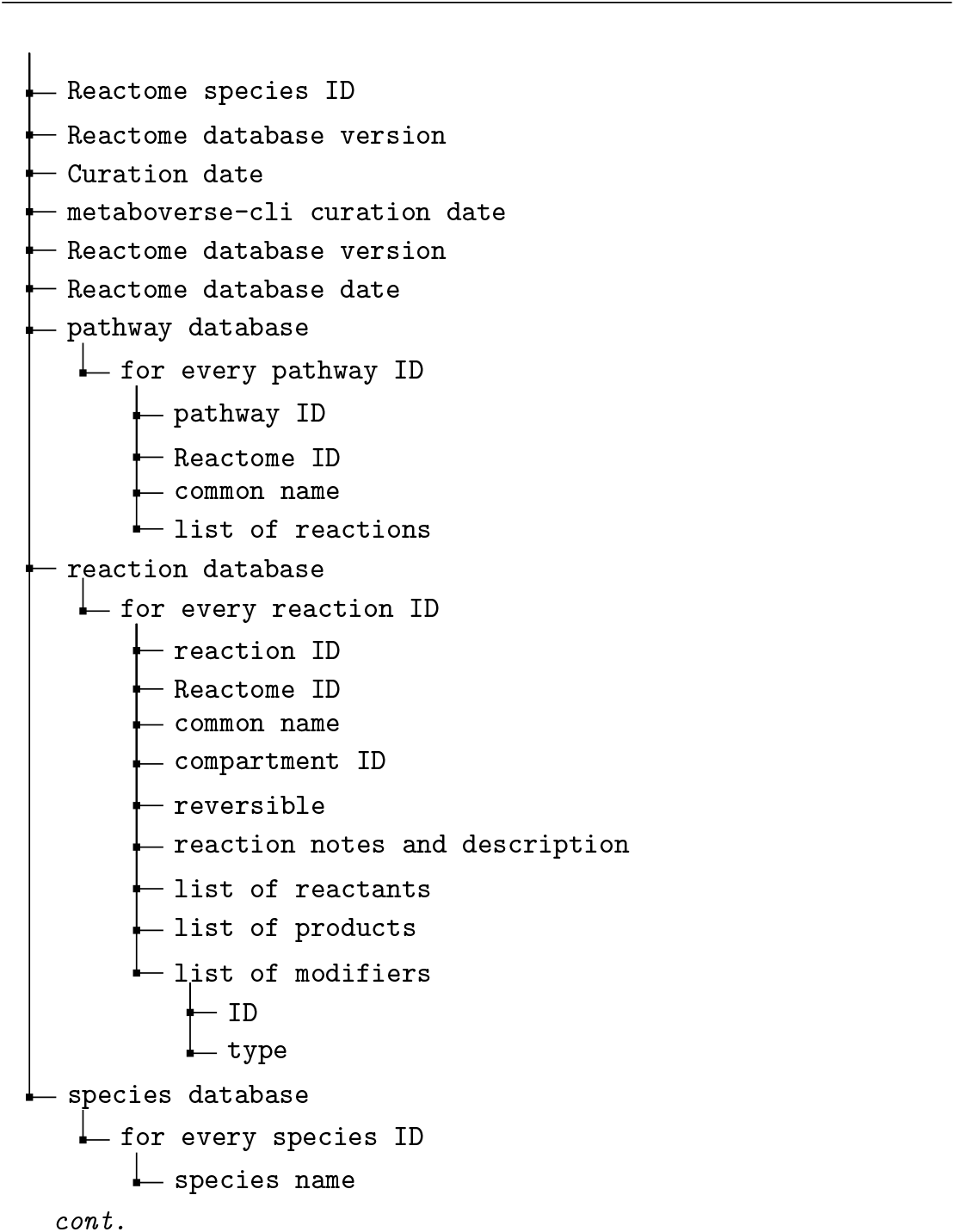
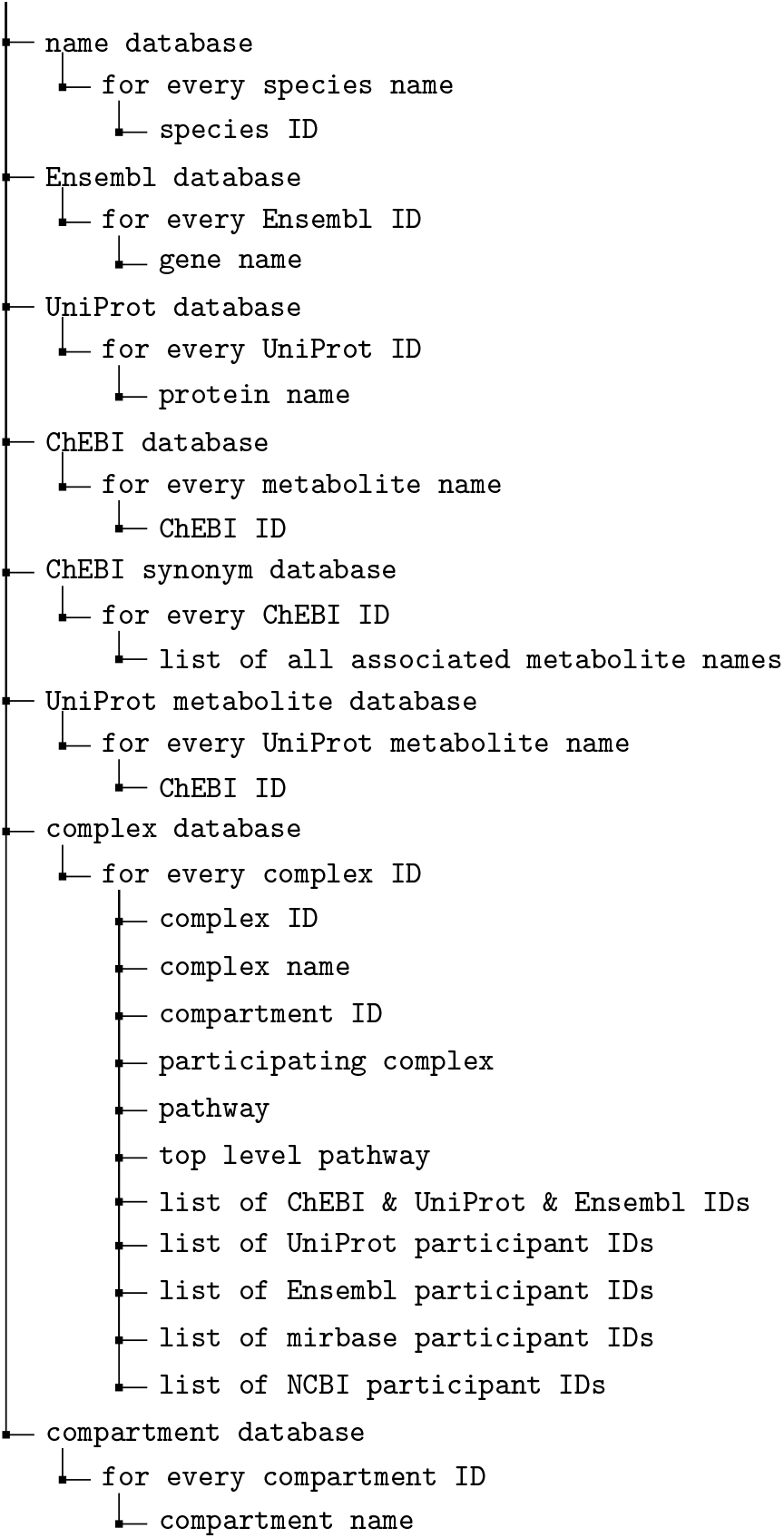
`mvdb` database structure.

After the relevant information is parsed from each table or record, the global network is propagated using the NetworkX framework [50] to generate nodes for each reaction and reaction component and edges connecting components to the appropriate reactions. In some cases, a separate ID is used to generate two nodes for the same metabolite within two separate compartments to aid in downstream visualization; however, user data for the given entity would be mapped to both nodes.

After the network is curated for the user-specified organism, each node’s degree (or magnitude of edges or connections) is determined to aid in the user’s downstream ability to avoid visualizing high-degree components, such as a proton or water, on the metabolic network, which can lead to visual network entanglement and cluttering and a decrease in computational performance [25]. The user may also choose to add metabolites or other components to a blocklist, which will lead to these entities being ignored during analysis and visualization. The resulting graph template is output as a JSON-formatted `.template.mvrs` file. A reaction neighbors JSON-formatted file is also output with the file suffix, `.nbdb`. These files, along with the initial `.mvdb` file are curated for each available organism with each release of Metaboverse to decrease back-end processing time on the user’s machine. These files are hosted at https://rutter.chpc.utah.edu/Metaboverse.

**Listing 2.**
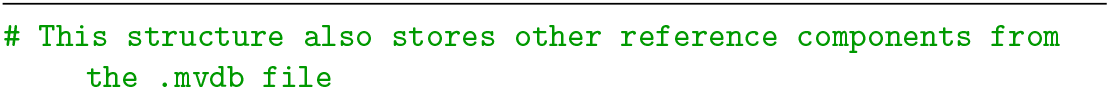
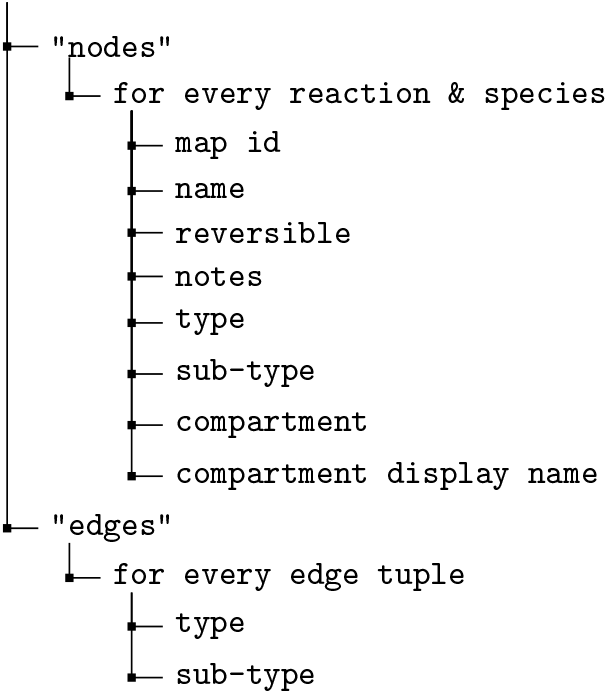
`template.mvrs` file structure.

**Listing 3.**
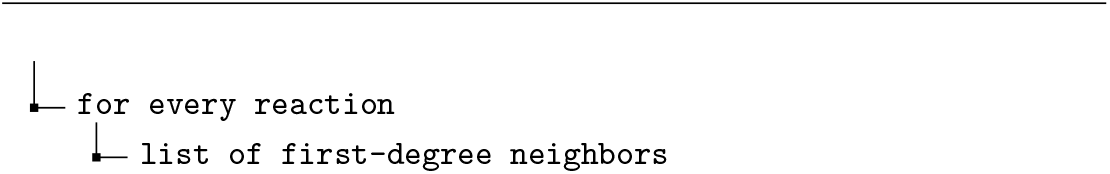
`nbdb` file structure.

**Listing 4.**
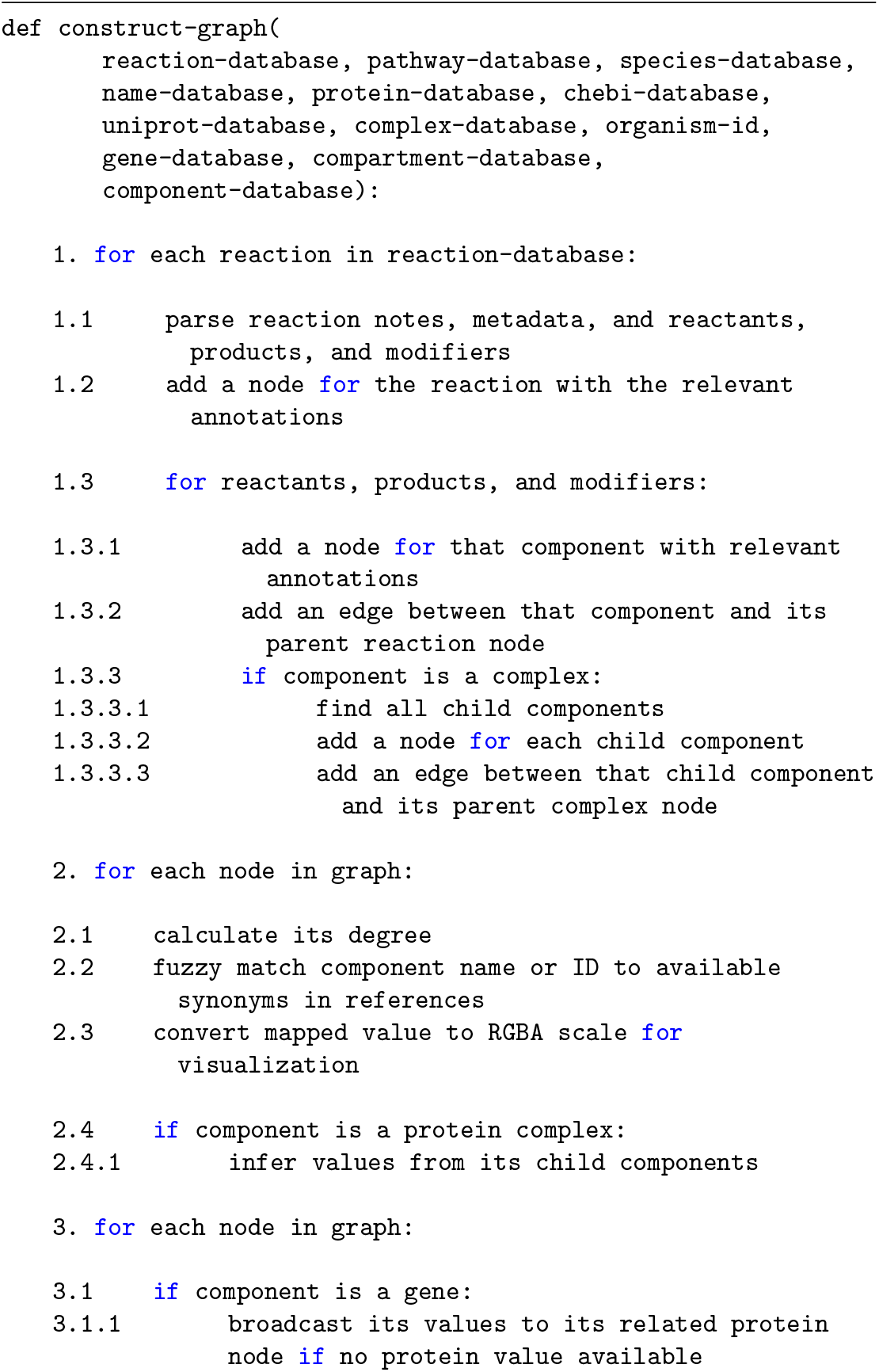
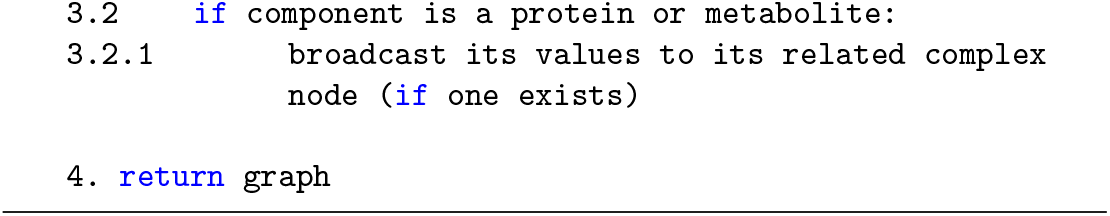
Graph construction overview.

In order to overlay user data on the global network, first, user-provided gene expression, protein abundance, and/or metabolite abundances’ names are mapped to Metaboverse compatible identifiers. For components that Metaboverse is unable to map, a table is returned to the user so they can provide alternative names to aid in mapping. Second, provided data values are mapped to the appropriate nodes in the network. In cases where gene expression data are available, but protein abundance values are missing, Metaboverse will take the average of the available gene expression values to broadcast to the protein node. For complexes, the median of all available component values (metabolites, proteins, etc.) are calculated in order to weight the inferred value towards values of more frequent magnitude. An aggregated p-value is inferred by multiplying the geometric mean of the p-values by *e*, as in [51, 52]. This method was chosen as it: 1) implies dependence between p-values, as can be expected between the regulation of components of a protein complex, 2) weights the resulting p-value towards significance and prevents penalizing the complex’s inferred p-value for one component with a poor p-value, and 3) ensures the resulting p-value is not lower than the minimum actual p-value from the set. Nodes for which values were inferred will be marked by a dashed border during visualization to clearly show which values are known and which were inferred. Statistical values are derived from the highest value of the components (assuming a scale of 1 denotes no statistical significance and 0 denotes high statistical significance).

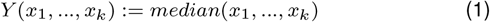

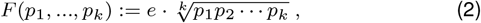

where *F* (*p*_1_, …, *p*_*k*_) ≤ 1.0.

### Collapsing reactions with missing expression or abundance values in user data

After data mapping is complete, Metaboverse will generate a collapsed network representation for optional viewing during later visualization. Metaboverse enforces a limit of up to three reactions that can be collapsed as data down a pathway should be inferred only so far. Reaction collapsing allows for partial matches between inputs and outputs of two reactions to account for key metabolic pathways where a metabolite that is output by one reaction may not be required for the subsequent reaction. For example, pperhaps ATP is produced by reaction A but is not required for reaction B. To perform a partial collapse, Metaboverse operates by largely the same scheme as outlined below, but additionally if a perfect match between reactions is not available, checks for partial matches by filtering out high-degree nodes (quartile 98 of all non-reaction node degrees) and then checking if, by default, at least 30% of the nodes match with its neighbor. Additional parameters for the reaction-collapse are as follows:

1. If a reaction has at least one known or inferred value for inputs (substrates) and one known or inferred value for outputs (products), the reaction will be left as is. During the entire reaction collapse step, known catalysts can be included when assessing whether a reaction has measured output values (increased catalyst should lead to more output in most cases), and inhibitors can be included when assessing whether the reaction has measured input values (increased inhibitor should lead to an accumulation of input in most cases). Catalysts and inhibitors are not included when determining reaction neighbors, as described below.
2. If a reaction has at least one known input, the input is left as is, and each reaction that shares the same inputs with the first reaction’s outputs is determined whether it has a measured output. If the neighbor reaction does not contain a known output value, the reaction is left as is. If the neighboring reaction does contain a measured output, the first reaction’s inputs and the neighboring reaction’s outputs are collapsed to form a single, pseudo-reaction between the two. If the reaction has at least one known output, the inverse is performed where neighbors with components identical to the reaction’s inputs are assessed for whether a collapsed reaction can be created.
3. If a reaction has no measured values, it is determined if the neighboring reactions on both sides (one sharing the reaction’s inputs and other sharing the reaction’s outputs) have measured values. If both neighbors contain a measured value, a collapsed pseudo-reaction is created, summarizing all three reactions.
4. All other reactions are maintained in the network.

For collapsed reactions, appropriate notes are included to describe the collapse. During visualization, these collapsed reactions are marked by black dashed edges and dashed node borders.

**Listing 5.**
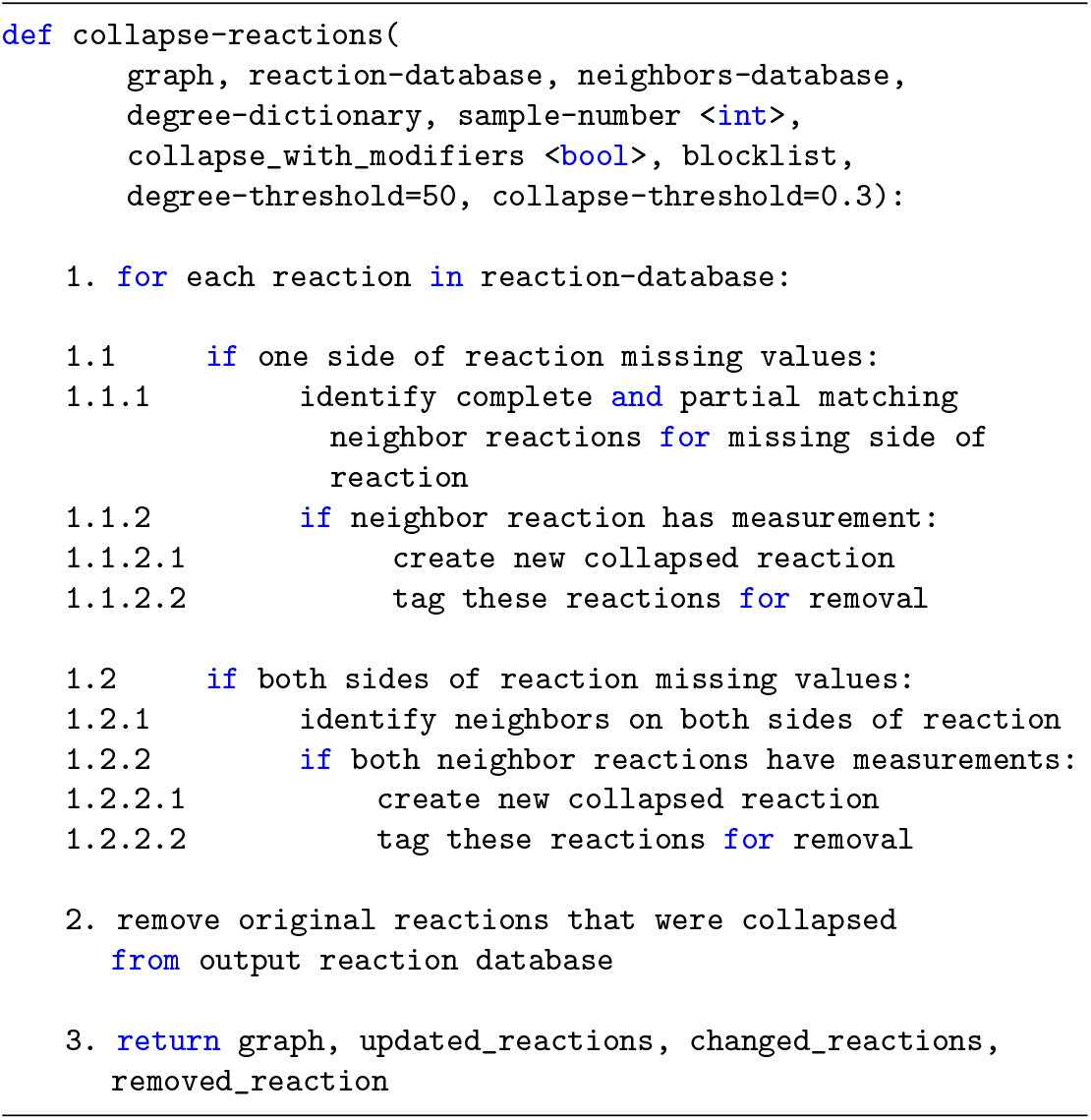
Reaction collapse schema.

### Regulatory pattern searches and sorting

Metaboverse provides a variety of different regulatory patterns for the user to explore. To identify a reaction-pattern is to compare some value that is computed from a reaction or a pathway with a user-specified threshold.

The identified reaction-patterns will be listed in a stamp view. Each stamp represents a reaction, with a glyph of the reaction, or the name of the pathway on it. In this stamp view, the identified patterns can be sorted according to three criteria: the number of pathways containing the reaction (not applicable for pathway pattern identification), the magnitude of the change of the computed value, and the statistical significance. When sorting by the number of pathways or the magnitude of the change, the identified reactions are arranged in order from the largest to the smallest. When sorting by the statistical significance, reactions with statistical significance on both the input side (substrates) and the output side (products) are listed first by the product of their maximum statistics, followed by the reactions with statistical significance on one of the two sides, and finally the reactions with no statistical significance on both sides. Within each tier, the reactions are sorted from lowest to highest p-values. For all values or statistics used in sorting, only those that determined the reaction-pattern are used.

**Listing 6.**
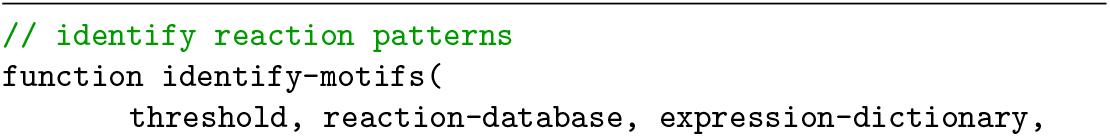
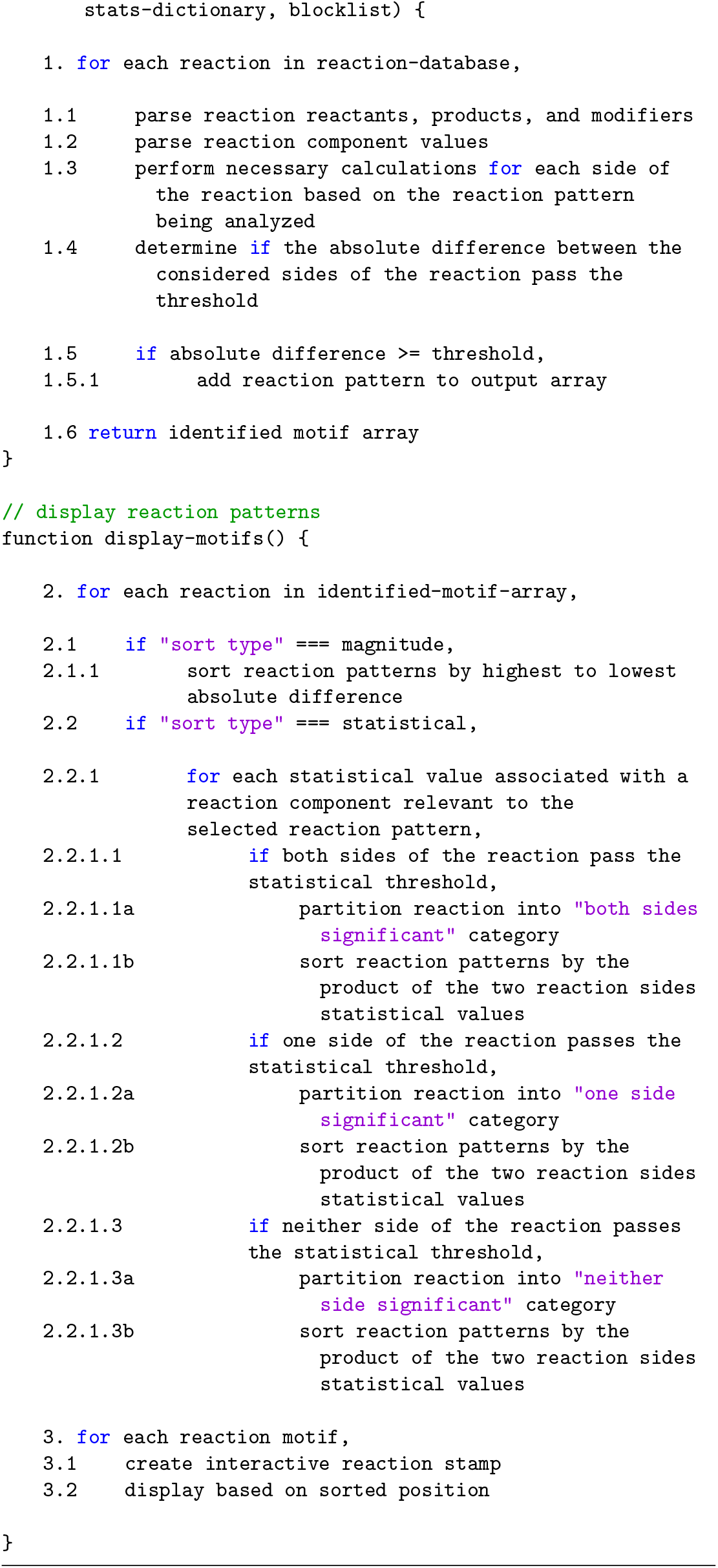
Pattern search schema.

When a reaction is selected from the stamp view, all the pathways containing the corresponding reactions will be listed below the stamp. Clicking on a pathway ID will draw the selected pathway in which the reaction-pattern was found, with all other reaction-patterns within this pathway also highlighted. For time-course and multi-condition datasets, the selected reaction-pattern’s total behavior is displayed below these windows as line-plots showing the reaction components’ behavior across all time-points or conditions.

### Nearest neighborhood searches and prioritization

To visualize all connections to a given network component, a user can select an entity (a gene, protein, or metabolite) and visualize all reactions in which the component is involved. By doing so, the user can visualize other downstream effects the change of one entity might have across the total network, which consequently aids in bridging and identifying any reaction that may occur between canonically annotated pathways. These neighborhoods can be expanded to view multiple downstream reaction steps and their accompanying genes, proteins, and metabolites by modulating the appropriate user option in the software.

The user can also limit which entities are shown by enforcing a degree threshold. By setting this value at 50, for example, the network would not show nodes that have 50 or more connections. One caveat, however, is that this feature will occasionally break synchronous pathways into multiple pieces if one of these high-degree nodes formed the bridge between two ends of a pathway.

### Perturbation networks

Perturbation networks are generated by searching each reaction in the total reaction network for any reaction where at least one component is significantly perturbed. The user can modify the necessary criteria to base the search on the expression or abundance value or the statistical value and can choose the thresholding value to be used. For the expression thresholding, the provided value is assumed to be the absolute value, so a thresholding value of 3 would include any reactions where at least one component showed a greater than 3 measured change or less than -3 measured change, the value of which is dependent on the data provided by the user. Thus, these networks could represent reactions where a component was perturbed to a significant degree on a log_2_ fold change scale, z-score scale, or other appropriate unit for that biological context. Once a list of perturbed reactions is collected, the network is constructed, including each of these reactions and their components. Perturbed neighboring reactions that share components are thus connected within the network, and perturbed reactions that are not next to other perturbed reactions are shown as disconnected sub-networks.

### Network visualization and exploration

Force-directed layouts of networks are constructed using D3 (https://d3js.org) by taking a user-selected pathway or entity and querying the reactions that are components of the selected pathway or entity. All inputs, outputs, modifiers, and other components of these reactions, along with edges where both source and target are found in the sub-network as nodes, are included and displayed. Relevant metadata, such as user-provided data and reaction descriptions, can be accessed by the user in real-time. To visualize a pathway, a user selects a pathway, and all component reactions and their substrates, products, modifiers, and metadata are queried from the total reaction database. Super-pathways help categorize these pathways and are defined as any pathway containing more than 200 nodes.

Time-course and multiple condition experiments are automatically detected from the user’s input data. When users provide these data and specifies the appropriate experimental parameters on the variable input page, they will have the option to provide time point or condition labels. Provided data should be listed in the data table in the same order that the labels are provided. Within all visualization modules, the data for each time point or condition can then be displayed using a slider bar, which will allow the user to cycle between time points or conditions.

Compartments are derived from Reactome annotations [5–7]. Compartment visualizations are generated using D3’s hull plotting feature. Compartment boundaries are defined at the reaction levels and made to encompass each reaction’s substrates, products, and modifiers for that given compartment.

Some performance optimization features are included by default to prevent computational overload. For example, nearest neighbor sub-networks with more than 1,500 nodes, or nodes with more than 500 edges, will not be plotted because the plotting of this information in real-time can be prohibitively slow.

### Packaging

The Metaboverse application is packaged using Electron (https://electronjs.org). Back-end network curation and data processing are performed using Python (https://www.python.org/) and the NetworkX [50], pandas [53, 54], NumPy [55], SciPy [54, 56], and Matplotlib (v3.4.2) [57] libraries. This back-end functionality is packaged as a single, operating system-specific executable using the PyInstaller library (https://www.pyinstaller.org) and is available to the app’s visual interface for data processing. Front-end visualization is performed using Javascript and relies on the D3 (https://d3js.org) and JQuery packages (https://jquery.com). Saving network representations to a PNG file is performed using the (https://github.com/edeno/d3-save-svg) and string-pixel-width (https://github.com/adambisek/string-pixel-width) packages. Documentation for Metaboverse is available at https://metaboverse.readthedocs.io. Continuous integration services are performed by GitHub Actions to routinely run test cases for each change made to the Metaboverse architecture. The Metaboverse source code can be accessed at https://github.com/Metaboverse/metaboverse. The code used to draft and revise this manuscript, as well as all associated scripts used to generate and visualize the data presented in this manuscript, can be accessed at https://github.com/Metaboverse/Metaboverse-manuscript/.

### Validation with biological data

#### Human lung adenocarcinoma metabolomics & analysis

Data were accessed from Metabolomics Workbench project PR000305 and processed as in our previous re-study of these data [25]. P-values were derived using a two-tailed, homoscedastic Student’s T-test and adjusted using the Benjamini-Hochberg correction procedure. Initial Kaplan-Meier survival analysis was performed using tools and data hosted on The Human Protein Atlas (v20.1; Released 2021-02-24) [29–31]. Survival analysis as displayed in this manuscript were performed in R (v4.0.3) using the survival (v3.2-11) and survminer (v0.4.9) packages. TCGA FPKM gene expression data were sourced from the Human Protein Atlas project (https://www.proteinatlas.org/download/rna_cancer_sample.tsv.zip) and clinical patient data were sourced from TCGA (https://portal.gdc.cancer.gov/projects/TCGA-LUAD). Clinical data were censored as “Dead” or “Alive” and “Alive” patients were right-censored using Days Since Last Follow-up. Patients were stratified into two gene expression groups (High, Low) using the optimized `surv cutpoint()` function from the survminer package (v0.4.9) with the minimum proportion for a group set at 0.2 [34].

#### Yeast strains

*Saccharomyces cerevisiae* BY4743 (MATa/α, his3/his3, leu2/leu2, ura3/ura3, met15/MET15, lys2/LYS2) was used to generate the *mct1*Δ. strain as described in [38].

#### Yeast growth assays

Growth assays were performed using S-min media with no uracil added and containing either 2% glucose or 2% raffinose. Equal numbers of wild-type or *mct1*Δ. yeast transformed with empty vector, *CTP1*-overexpression, and *CTP1*-C-terminal GFP-overexpression plasmids were spotted as 10-fold serial dilutions during mid-log phase (OD_600_=0.3-0.6). Plates were incubated at 30 °C for 2–3 days before imaging.

#### RNA-sequencing sample preparation and analysis

RNA sequencing data were generated by growing *Saccharomyces cerevisiae* biological replicates for strains *mct1*Δ (n=4) and wild-type (n=4). Briefly, cells were grown in glucose and switched to raffinose-supplemented growth medium for 0, 3, and 12 hours such that at the time of harvest, cultures were at OD_600_=1. Cultures were flash-frozen, and later total RNA was isolated using the Direct-zol kit (Zymo Research) with on-column DNase digestion and water elution. Sequencing libraries were prepared by purifying intact poly(A) RNA from total RNA samples (100-500 ng) with oligo(dT) magnetic beads, and stranded mRNA sequencing libraries were prepared as described using the Illumina TruSeq Stranded mRNA Library Preparation Kit (RS-122-2101, RS-122-2102). Purified libraries were qualified on an Agilent Technologies 2200 TapeStation using a D1000 ScreenTape assay (cat# 5067-5582 and 5067-5583). The molarity of adapter-modified molecules was defined by quantitative PCR using the Kapa Biosystems Kapa Library Quant Kit (cat# KK4824). Individual libraries were normalized to 5 nM, and equal volumes were pooled in preparation for Illumina sequence analysis. Sequencing libraries (25 pM) were chemically denatured and applied to an Illumina HiSeq v4 single-read flow cell using an Illumina cBot. Hybridized molecules were clonally amplified and annealed to sequencing primers with reagents from an Illumina HiSeq SR Cluster Kit v4-cBot (GD-401-4001). Following transfer of the flowcell to an Illumina HiSeq 2500 instrument (HCSv2.2.38 and RTA v1.18.61), a 50 cycle single-read sequence run was performed using HiSeq SBS Kit v4 sequencing reagents (FC-401-4002).

Sequence FASTQ files were processed using XPRESSpipe (v0.6.0) [58]. Batch and log files are available at https://github.com/Metaboverse/Metaboverse-manuscript/tree/main/data/sce_mct1_omics. Notably, reads were trimmed of adapters (AGATCGGAAGAGCACACGTCTGAACTCCAGTCA). Based on library complexity quality control, de-duplicated alignments were used for read quantification due to the high number of duplicated sequences in each library. Differential expression analysis was performed using DESeq2 [59] by comparing *mct1*Δ. samples with wild-type samples at the 12-hour time-point to match the steady-state proteomics data. log_2_(fold change) and false discovery rate (“p-adj”) values were extracted from the DESeq2 output.

#### Proteomics analysis

Steady-state quantitative proteomics data previously processed and obtained from [38]. Briefly, cells were grown in glucose and switched to raffinose-supplemented growth medium overnight and harvested at mid-log phase. For this analysis, we compared the *mct1*Δ. (n=3) with the wild-type (n=3) cell populations. log_2_(fold change) values and Benjamini-Hochberg corrected p-values were generated by comparing *mct1*Δ. with the wild-type cells. P-values were generated before correction using a 2-tailed, homoscedastic Student’s T-test.

#### Gas chromatography metabolomics sample preparation and metabolite extraction

Metabolomics data were generated by growing the appropriate yeast strains in synthetic complete media supplemented with 2% glucose until they reached saturation (n=6; except in one 3-hour wild-type sample, where n=5). Cells were then transferred to S-min media containing 2% raffinose and leucine and harvested after 0, 15, 30, 60, and 180 minutes (n=6/time-point/strain, except for the 3-hour wild-type samples, where n=5) at OD_600_=0.6-0.8.

A 75% boiling ethanol (EtOH) solution containing the internal standard d4-succinic acid (Sigma 293075) was then added to each sample. Boiling samples were vortexed and incubated at 90 °C for 5 minutes. Samples were then incubated at -20 °C for 1 hour. After incubation, samples were centrifuged at 5,000 x g for 10 minutes at 4°C. The supernatant was then transferred from each sample tube into a labeled, fresh 13×100mm glass culture tube. A second standard was then added (d27-myristic acid CDN Isotopes: D-1711). Pooled quality control samples were made by removing a fraction of collected supernatant from each sample, and process blanks were made using only extraction solvent and no cell culture. The samples were then dried *en vacuo*. This process was completed in three separate batches.

#### Gas chromatography mass spectrometry analysis of samples

All GC-MS analysis was performed with an Agilent 5977b GC-MS MSD-HES and an Agilent 7693A automatic liquid sampler. Dried samples were suspended in 40 *µ*L of a 40 mg/mL O-methoxylamine hydrochloride (MOX) (MP Bio #155405) in dry pyridine (EMD Millipore #PX2012-7) and incubated for 1 hour at 37 °C in a sand bath. 25 *µ*L of this solution were added to auto sampler vials. 60 *µ*L of N-methyl-N-trimethylsilyltrifluoracetamide (MSTFA with 1%TMCS, Thermo #TS48913) were added automatically via the auto sampler and incubated for 30 minutes at 37 °C. After incubation, samples were vortexed, and 1 *µ*L of the prepared sample was injected into the gas chromatograph inlet in the split mode with the inlet temperature held at 250 °C. A 10:1 split ratio was used for the analysis of the majority of metabolites. For those metabolites that saturated the instrument at the 10:1 split concentration, a split of 50:1 was used for the analysis. The gas chromatograph had an initial temperature of 60 °C for 1 minute followed by a 10 °C/min ramp to 325 °C and a hold time of 5 minutes. A 30-meter Phenomenex Zebron AB-5HT with 5m inert Guardian capillary column was employed for chromatographic separation. Helium was used as the carrier gas at a rate of 1 mL/min.

Data were collected using MassHunter software (Agilent). Metabolites were identified, and their peak area was recorded using MassHunter Quant. These data were transferred to an Excel spreadsheet (Microsoft, Redmond WA). Metabolite identity was established using a combination of an in-house metabolite library developed using pure purchased standards, the NIST (https://www.nist.gov) and Fiehn libraries [60]. Resulting data from all samples were normalized to the internal standard d4-succinate. P-values were derived using a homoscedastic, two-tailed Student’s T-test and adjusted using the Benjamini-Hochberg correction procedure.

#### Liquid chromatography metabolomics sample preparation and metabolite extraction

Metabolomics data were generated by growing the appropriate yeast strains in synthetic complete media supplemented with 2% glucose until they reached saturation (n=3). Cells were then transferred to S-min media containing 2% raffinose and leucine and harvested after 0, 15, 30, 60, and 180 minutes (n=6/time-point/strain, except for the 3-hour wild-type samples, where n=5) at OD_600_=0.6-0.8.

The procedures for metabolite extraction were performed as previously described [61]. Yeast cultures were pelleted, snap frozen and kept at *−* 80°C. 5ml of 75% boiled ethanol was added to every frozen pellet. Pellets were vortexed and incubated at 90°C for 5 minutes. All samples were then centrifuged at 5,000 Relative Centrifugal Force (RCF) for 10 minutes. Supernatants were transferred to fresh tubes, evaporated overnight in a Speed Vacuum, and then stored at *−*80°C until they were run on the mass spectrometer.

#### Liquid chromatography mass spectrometry analysis of samples

The conditions for liquid chromatography are described in previous studies [62, 63]. Briefly, a hydrophilic interaction liquid chromatography method (HILIC) with an Xbridge amide column (100 *×* 2.1 mm, 3.5 µm) (Waters) was employed on a Dionex (Ultimate 3000 UHPLC) for compound separation and detection at room temperature. The mobile phase A was 20 mM ammonium acetate and 15 mM ammonium hydroxide in water with 3% acetonitrile, pH 9.0, and the mobile phase B was acetonitrile. The linear gradient was as follows: 0 min, 85% B; 1.5 min, 85% B, 5.5 min, 35% B; 10 min, 35% B, 10.5 min, 35% B, 14.5 min, 35% B, 15 min, 85% B, and 20 min, 85% B. The flow rate was 0.15 ml/min from 0 to 10 min and 15 to 20 min, and 0.3 ml/min from 10.5 to 14.5 min. All solvents were LCMS grade and purchased from Thermo Fisher Scientific.

Mass spectrometry was performed as described in previous studies [62, 63]. Briefly, the Q Exactive MS (Thermo Scientific) is equipped with a heated electrospray ionization probe (HESI), and the relevant parameters are as listed: evaporation temperature, 120°C; sheath gas, 30; auxiliary gas, 10; sweep gas, 3; spray voltage, 3.6 kV for positive mode and 2.5 kV for negative mode. Capillary temperature was set at 320°C, and S-lens was 55. A full scan range from 60 to 900 (m/z) was used. The resolution was set at 70,000. The maximum injection time was 200 ms. Automated gain control (AGC) was targeted at 3,000,000 ions.

Data were collected, metabolites were identified, and their peak area was recorded using El-MAVEN software [64–66]. These data were transferred to an Excel spreadsheet (Microsoft, Redmond WA). Metabolite identity was established using a combination of an in-house metabolite library developed using pure purchased standards, the NIST (https://www.nist.gov) and Fiehn libraries [60]. Resulting data from all samples were normalized to the internal standard d4-succinate. P-values were derived using a homoscedastic, two-tailed Student’s T-test and adjusted using the Benjamini-Hochberg correction procedure.

#### Correlation analysis

To correct the expression bias arising from highly expressed genes, gene expression data were first corrected using spatial quantile normalization (SpQN; v1.0.0) for each dataset with the first four principle components being removed for each dataset [45]. Genes were considered co-expressed in refine.bio datasets if r *>* 0.5 and in the lab-generated wild-type data if *>* 0.75.

Gene ontology enrichment analysis was performed by processing the correlated gene sets from each dataset using the PANTHER Overrepresentation Test (v16; Released 20210224) on the GO biological process complete annotation dataset (https://doi.org/10.5281/zenodo.4735677; Released 2021-05-01) [67, 68] via the Gene Ontology (GO) Resource [69, 70]. Enrichments were determined using Fisher’s Exact Test and p-values were corrected using the PANTHER false discovery rate calculation [67, 68]. Enrichments were prioritized by fold change. For overlapping GO terms, the GO term was the highest fold change was used for the visualization. Enrichment FDRs and fold changes were visualized as bubble plots generated using seaborn (v0.11.0) and Matplotlib (v3.4.2) [57, 71]. Scatterplots of co-expressed genes against the gene of interest were generated using the `regplot()` function from seaborn (v0.11.0) and Matplotlib (v3.4.2) [57, 71].

#### Heatmap and boxplot visualization

Heatmaps were generated using the `clustermap()` function from seaborn (v0.11.0) and Matplotlib (v3.4.2) using custom gene, protein, or metabolite lists [57, 71]. Heatmap values were mean-centered at 0 (z-score). Hierarchical clustering was performed where indicated by the linkage lines using a simple agglomerative (bottom-up) hierarchical clustering method (or UPGMA (unweighted pair group method with arithmetic mean)). Gene comparison boxplots were generated using the `swarmplot()` and `boxplot()` functions from seaborn (v0.11.0) and Matplotlib (v3.4.2) [57, 71].

## Data availability

Gene expression counts for lung adenocarcinomas were sourced from the Human Protein Atlas project’s TCGA FPKM gene expression data (https://www.proteinatlas.org/download/rna_cancer_sample.tsv.zip) and clinical patient data were sourced from TCGA (https://portal.gdc.cancer.gov/projects/TCGA-LUAD).

*mct1*Δ. and accompanying wild-type transcriptomics time-course data are deposited at the GEO repository under identifier GSE151606. Metabolomics data are deposited at the Metabolomics Workbench repository under project identifier PR000961, study identifier ST001401 (DOI: 10.21228/M8GD71).

For gene co-expression analyses, all yeast samples available in refine.bio were accessed on March 16, 2021 [44].

The curated networks for these data are available at https://github.com/Metaboverse/Metaboverse-manuscript/tree/main/manuscript_files. Networks were generated by taking the 12-hour transcriptomics and proteomics datasets with their appropriate log_2_(fold change) and statistical values, along with the 0, 15, 30, 60, and 180 minute metabolomics datasets with their respective log_2_(fold change) and statistical values and layering these data on the *Saccharomyces cereviseae* global reaction network as curated by Metaboverse from the Reactome database [5–7]. reaction-patterns and perturbation network analyses were performed within the Metaboverse platform.

## Code availability

The Metaboverse source code is available at https://github.com/Metaboverse/Metaboverse and https://github.com/Metaboverse/metaboverse-cli. The latest version of the software can be found at https://github.com/Metaboverse/Metaboverse/releases/latest. The source code and data for this manuscript and the subsequent analyses are available at https://github.com/Metaboverse/Metaboverse-manuscript/.

## Acknowledgments

We thank Thomas O’Connell, Pauline Trébulle, and Joshua Rabinowitz for valuable feedback and feature suggestions during beta-testing of the software. We thank Alex J. Bott, Kevin G. Hicks, Jeffrey T. Morgan, and other members of the Rutter lab for their thoughtful insights and suggestions. We thank Geanette Lam for assistance in generating the *CTP1*-GFP fusion plasmid used in this study. We thank Brian Dalley and the University of Utah High-Throughput Genomics Core for help with RNA library preparation and sequencing. We thank Sarah Fogarty and Jacob M. Winter for critical feedback during drafting of the manuscript. We also thank Christine Pickett for providing editing services for this manuscript. The support and resources from the Center for High-Performance Computing at the University of Utah are gratefully acknowledged.

J.A.B. is supported by the National Cancer Institute (NCI) of the National Institutes of Health (NIH) under award number F99CA253744 (https://www.cancer.gov) and received additional support from the National Institute of Diabetes and Digestive and Kidney Diseases (NIDDK) Inter-disciplinary Training Grant T32 Program in Computational Approaches to Diabetes and Metabolism Research, 1T32DK11096601 to Wendy W. Chapman and Simon J. Fisher (https://www.niddk.nih.gov). S.M.N. received support from The United Mitochondrial Disease Foundation PF-15-046 (https://www.umdf.org) and the American Cancer Society PF-18-106-01 (https://www.cancer.org) postdoctoral fellowships, along with T32HL007576. J.E.C. is funded by S10OD016232, S10OD021505, and U54DK110858. This work was supported by funds from the NIDDK fellowship 1T32DK11096601 and the NCI fellowship 1F99CA253744 (to J.A.B.) (https://www.niddk.nih.gov, https://www.cancer.gov), the NSF grants DBI-1661375 and IIS-1513616 (to B.W.) (https://www.nsf.gov), and the NIH grant R35GM131854 (to J.R.) (https://www.nih.gov). The computational resources used were partially funded by the NIH Shared Instrumentation Grant 1S10OD021644-01A1 (https://www.nih.gov). Mass spectrometry equipment was obtained through NCRR Shared Instrumentation Grant 1S10OD016232-01, 1S10OD018210-01A1, and 1S10OD021505-01 (to J.E.C.). The funders had no role in study design, data collection and analysis, decision to publish, or preparation of the manuscript. The content of this manuscript is solely the responsibility of the authors and does not necessarily represent the official views of the National Institutes of Health (NIH).

## Author contributions

Y.Z. and Y.O. contributed equally to this work. Conceptualization: J.A.B., T.C.W., B.W., J.R.; Supervision: J.A.B., B.W., J.R.; Project Administration: J.A.B.; Investigation: J.A.B., Y.O., T.C.W., A.A.C., M.E.C., S.M.N., T.V.; Formal Analysis: J.A.B., Y.Z., Y.O., A.A.C., T.V.; Software: J.A.B., Y.Z., I.G.; Methodology: J.A.B., Y.Z.; Validation: J.A.B., Y.O., A.A.C., M.E.C., I.G.; Data Curation: J.A.B., Y.O., T.C.W., A.A.C., S.M.N., T.V.; Resources: J.A.B., J.E.C., B.W., J.R.; Funding Acquisition: J.A.B., J.E.C., B.W., J.R.; Writing - Original Draft Preparation: J.A.B.; Writing - Review & Editing: J.A.B., Y.Z., Y.O., T.C.W., A.A.C., T.V., I.G., S.N.M., J.E.C., B.W., J.R.; Visualization: J.A.B., Y.Z., Y.O.

## Supplemental Data & Figures

### S1 Text: A brief overview of representative metabolic network analysis tools

We will highlight four such representative and popular tools for their respective properties, though many more exist [1]. First is MetaboAnalyst, which relies heavily on set enrichment methods for the analysis of data or examining the belongingness of sets of significantly changed analytes (i.e., metabolite, protein, or gene measurements), for extracting interesting information. Network visualization is available, but it focuses primarily on interaction networks, and its ability to extract regulatory information is limited, particularly in an automated fashion [2,3]. Second is Cytoscape, which serves as a general go-to platform for representing and visualizing biological or other networks. One strength of Cytoscape is the ability to design apps or plug-ins to conduct customized analyses; however, comprehensive and metabolism-specific regulatory identification methods are unavailable [4]. One plug-in for Cytoscape that focuses on metabolic data is MetScape, but again, this tool is generally limited to pathway enrichment, correlation networks, and data visualization and does not integrate approaches to identify regulatory mechanisms at the reaction-level within the data [5-7]. MetExplore focuses on the curation of networks and is particularly useful for collaborative annotation of emerging models of organisms with incomplete metabolic network curations. It additionally can layer experimental data on the network for visualization [8,9]. A companion tool to MetExplore is MetExploreViz, which enables interactive and flexible visualization of omics data on metabolic networks [10]. Reactome, which our tool uses for the curation of biological networks, also offers analytical tools for user data, but again relies on set enrichment or manual methods for identifying patterns [11-13]. While all have their respective utility, there is a pressing need for tools that integrate these features and automate pattern and trend detection across metabolic networks to extract regulatory and other features from data. For an in-depth discussion of other metabolomics and metabolism computational tools, we refer readers to the resource paper, [14].

---

**S1 Fig:**
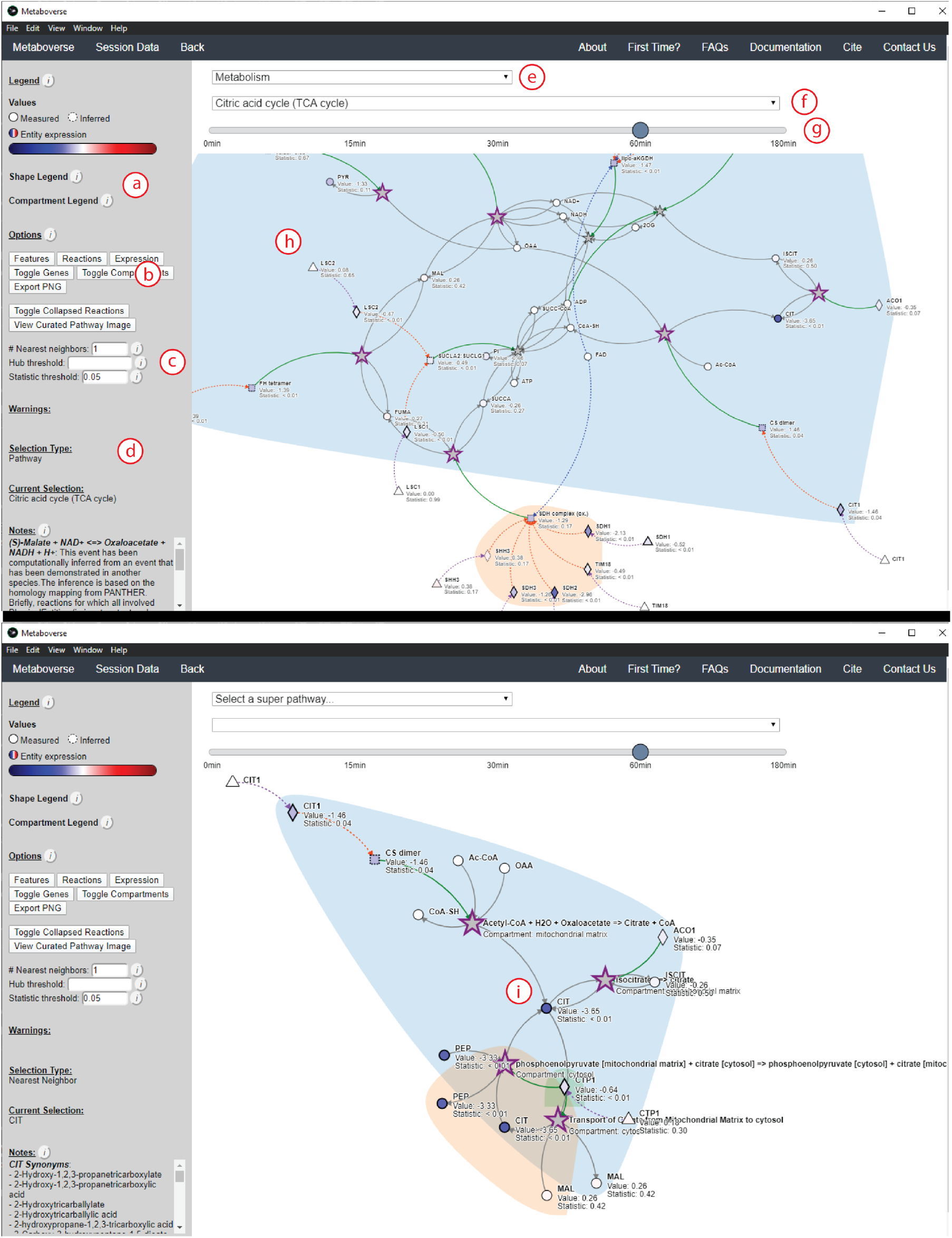
Example of the General Exploration module. **a**. By hovering on the icon, users can view the shape legend. Below the shape legend is the compartment legend, where all currently displayed compartments in the selected pathway will be shown. **b**. Users can toggle features on or off using the appropriate buttons. For example, users could choose to display all reaction names, not just when a reaction is selected. **c**. Users can modulate graphing parameters, such as the number of reaction neighborhoods to graph when selecting a network component for nearest neighborhood analysis. **d**. Notes about the selected reaction or synonyms for the selected reaction component appear here, as well as other information. **e**. The drop-down menu for the selection of a super-pathway. **f**. The drop-down menu for the selection of a specific pathway within the selected super-pathway. **g**. For time-course and multi-condition datasets, a slider bar will appear that users can use to change value shading of nodes and highlighted reaction-patterns. **h**. Viewing area for the selected pathway. **i**. When users double-click on a reaction component, a nearest neighborhood graph will be displayed. In this instance, citrate was selected, and all reactions citrate is involved in across all pathways are plotted.

**S2 Fig:**
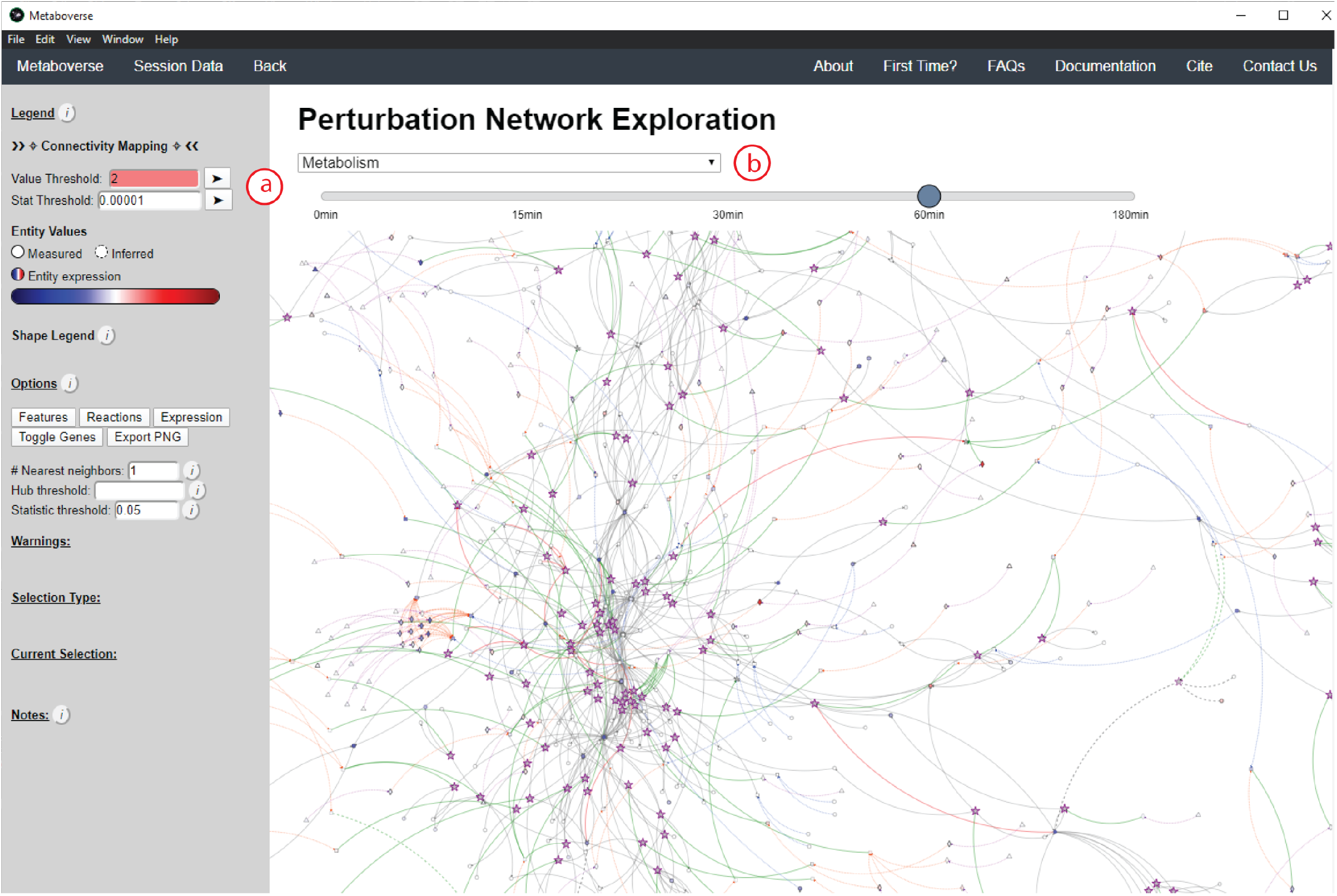
Example of the Perturbation Networks module. **a**. Users can select what constitutes a perturbed reaction by modulating the appropriate thresholds. Only one type of perturbation (by magnitude or by statistical value) will be used. **b**. Users select a super-pathway, for which all perturbed reactions belonging to that super-pathway are selected and displayed. Reactions that are neighbors and perturbed will be shown stitched together within the network, even if these reactions are connected across different pathways.

**S3 Fig:**
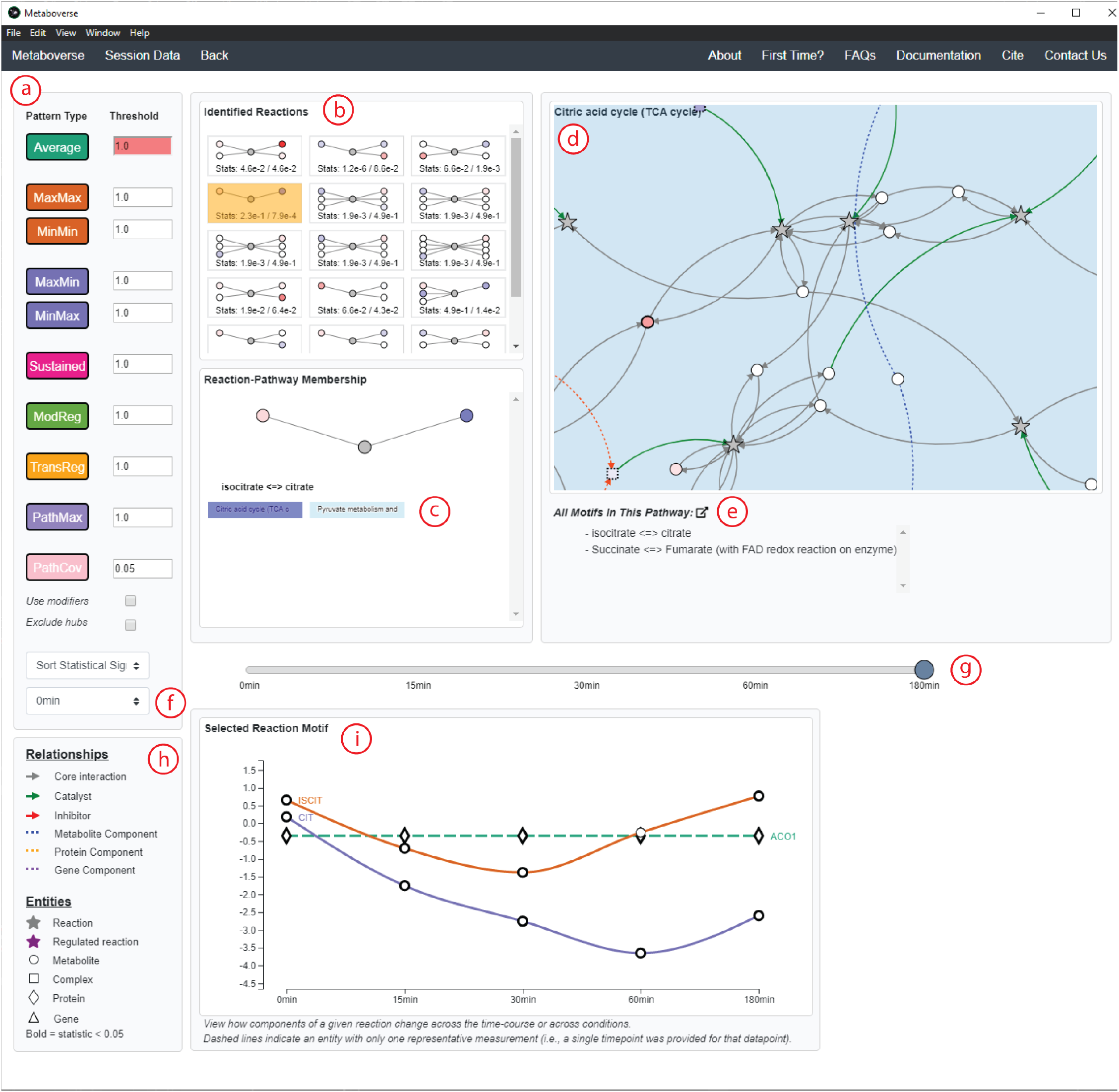
Example of the Pattern Analysis module. **a**. The patterns currently available to search in Metaboverse are displayed in this panel. Users can adjust the thresholds for the given pattern using the appropriate fields. **b**. For the selected pattern type, all available reaction-patterns within the organism’s network will appear here. Reactions will be sorted by statistical values, as provided by the user, that were relevant to the identified pattern. Green stamps indicate both sides of the reaction contained significant values, yellow stamps indicate one side of the reaction contained significant values, and gray stamps indicate neither side of the reaction contained significant values. **c**. This panel displays all of the pathways in which the reaction-pattern can be found. The graphic provides a simplified view of the reaction’s primary substrates and products, with the nodes shaded by associated value. A user can select one of these pathways for viewing. **d**. The selected pathway is displayed here, with all reaction-patterns in the pathway highlighted with a bold purple border. **e**. If a user would like to view the selected pathway within the ‘Explore’ module, they can click the icon to open the pathway in a new window. All reaction-patterns within the displayed pathway are listed below. **f**. For time-course or multi-condition datasets, the user can choose to display reaction-patterns for the selected time-point or condition that are not present in another time-point or condition. **g**. For time-course or multi-condition datasets, a slider bar will appear with the time-points or conditions for the user to select. **h**. This panel provides a shape legend for the different shape types used in Metaboverse. **i**. For time-course or multi-condition datasets, a line plot will appear for the selected reaction. While panels above only display the behavior of that reaction-pattern at a single time-point or condition, the line plot will display that reaction’s behavior across all of the available time-points or conditions.

**S4 Fig:**
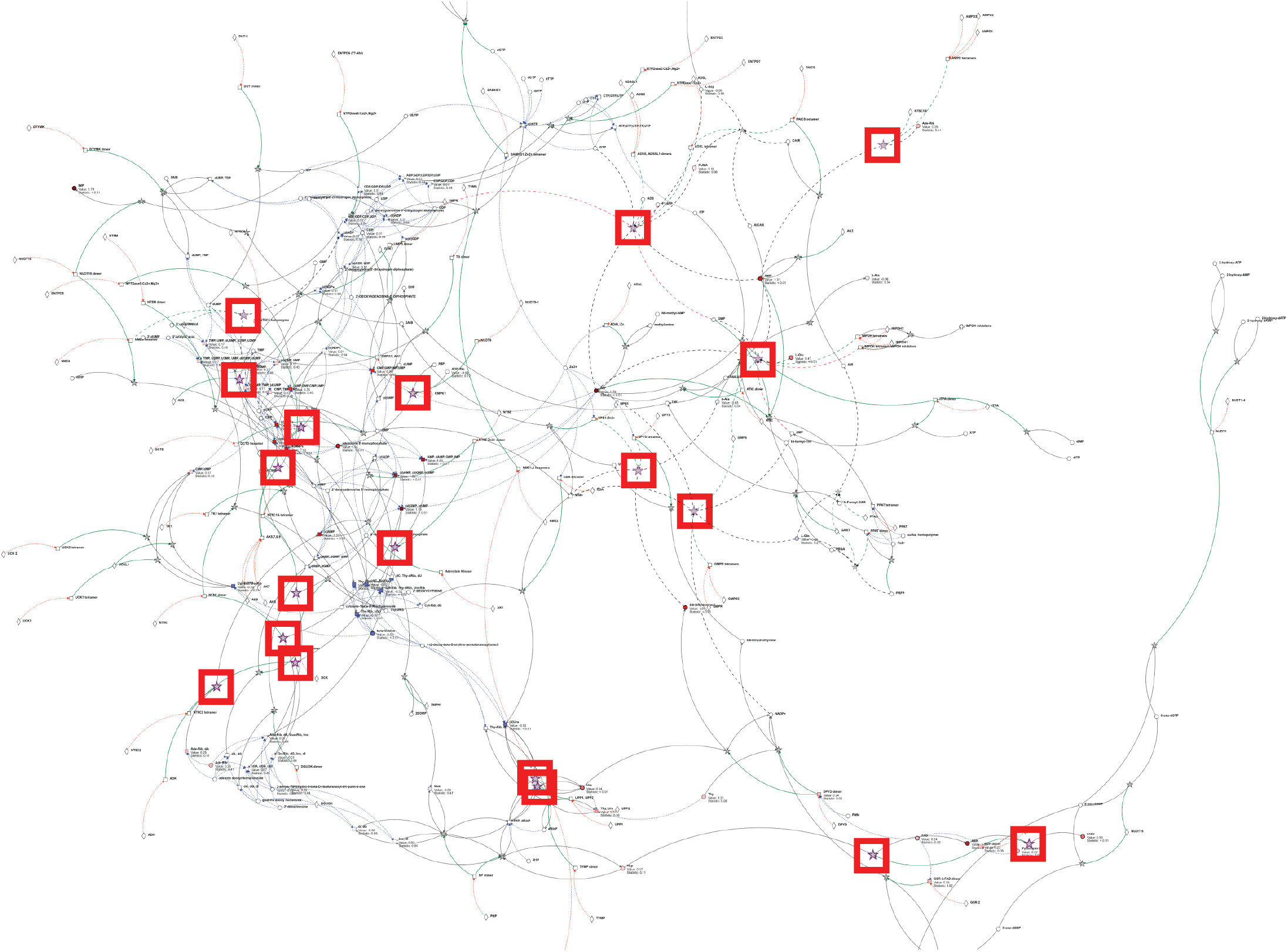
Metaboverse identifies reaction patterns in nucleotide metabolism. Reaction patterns (red boxes) in metabolism of nucleotides pathway (R-HSA-15869).

**S5 Fig:**
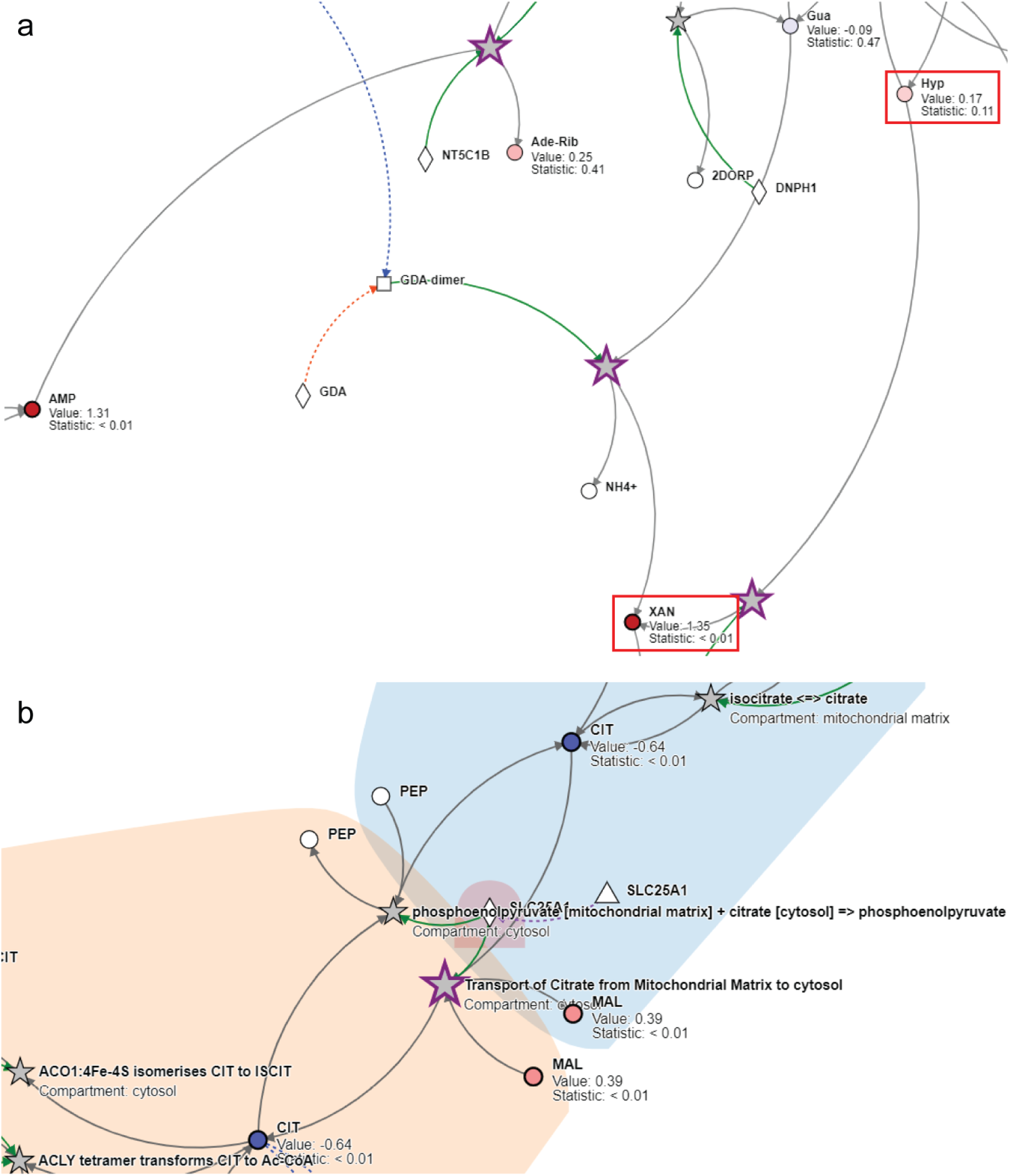
Metaboverse identifies reaction patterns in xanthine and TCA metabolism. **a**. Identification of xanthine regulation by both the pattern recognition and perturbation analysis modules. **b**. Disruptions of TCA metabolism support canonical disruptions during adenocarcinoma development. Metabolomics values are shown as node shading, where an increasingly blue shade indicates downregulation, and an increasingly red shade indicates upregulation. Measured log_2_(fold change) and statistical values for each entity are displayed below the node name. A gray node indicates a reaction. A bold gray node with a purple border indicates a motif at this reaction. Circles indicate metabolites, squares indicate complexes, and diamonds indicate proteins. Gray edges indicate core relationships between reaction inputs and outputs. Green edges indicate a catalyst, and red edges indicate inhibitors. Dashed blue edges point from a metabolite component to the protein complex in which it is involved. Dashed orange edges point from a protein component to the protein complex in which it is involved. Protein complexes with dashed borders indicate that the values displayed on that node were inferred from the constituent protein and metabolite measurements. Hub limit was set at 30 during generation of the network visualization as shown in sub-panel **b**.

**S6 Fig:**
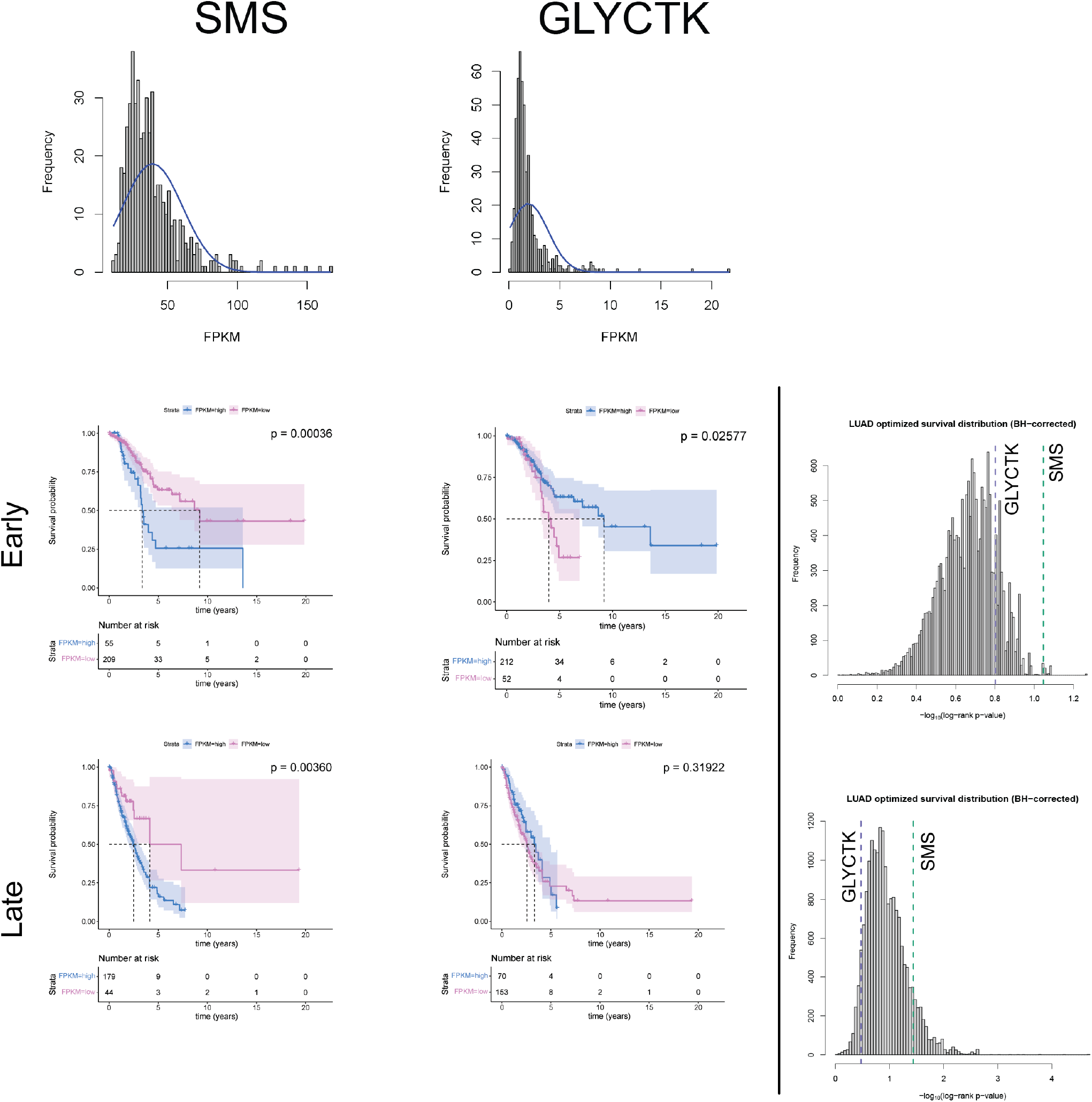
Overall survival outcomes correlations of SMS and GLYCTK gene expression in early stage adenocarinomas are stronger than in later stage adenocarcinomas. (top) gene FPKM distributions for SMS and GLYCTK. (middle) Kaplan-Meier plots for early stage (stage IA-B) samples for SMS and GLYCTK and distribution of all genes’ Benjamini-Hochberg log-rank p-values. (bottom) Kaplan-Meier plots for late stage (stage II+) samples for SMS and GLYCTK and distribution of all genes’ Benjamini-Hochberg log-rank p-values. Shading in Kaplain-Meier plots indicates 95% confidence intervals for each expression group. Dashed lines indicate median survival times for each expression group. Risk tables are displayed below each Kaplan-Meier plot, and include the number of individuals in each risk category at time = 0 years.

**S7 Fig:**
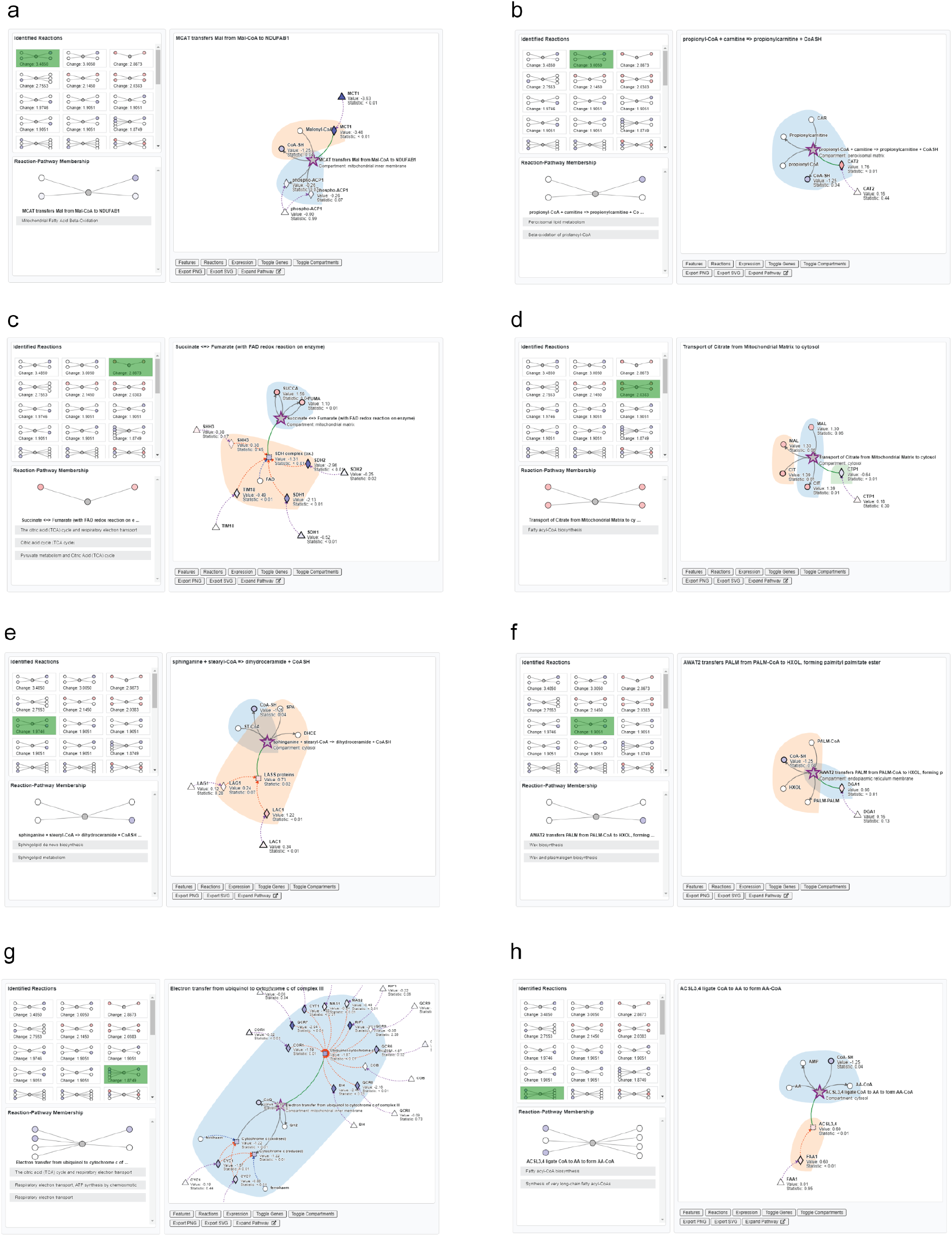
Top-ranking ModReg reaction patterns identified in steady-state proteomics and metabolomics data in the *mct1*Δ background. Stamp view snapshots of the eight top-ranking `ModReg` reaction patterns in the *mct1*Δ vs. wild-type comparison using steady-state (12 hr) proteomics and metabolomics data. Reaction patterns were sorted by difference in magnitude of the different relevant components. Only results where the input/output and modifier were both statistically significant are shown. RNA-sequencing comparisons contained n=4 in each group, proteomics comparisons contained n = 3 in each group, and metabolomics comparisons contained n = 6 in each each comparison group, except for the 3-hour wild-type group, which contained n = 5. **a**. “MCAT transfers Mal from Mal-CoA to NDUFAB1”. **b**. “propionyl-CoA + carnitine =*>* propionylcarnitine + CoA SH”. **c**. “Succinate *<*=*>* Fumarate (with FAD redox reaction on enzyme)”. **d**. “Transport of Citrate from Mitochondrial Matrix to cytosol”. **e**. “sphinganine + stearyl-CoA =*>* dihydroceramide + CoASH”. **f**. “AWAT2 transfers PALM from PALM-CoA to HXOL, forming palmityl palmitate ester”. **g**. “Electron transfer from ubiquinol to cytochrome c of complex III”. **h**. “ACSL3,4 ligate CoA to AA to form AA-CoA”.

**S8 Fig:**
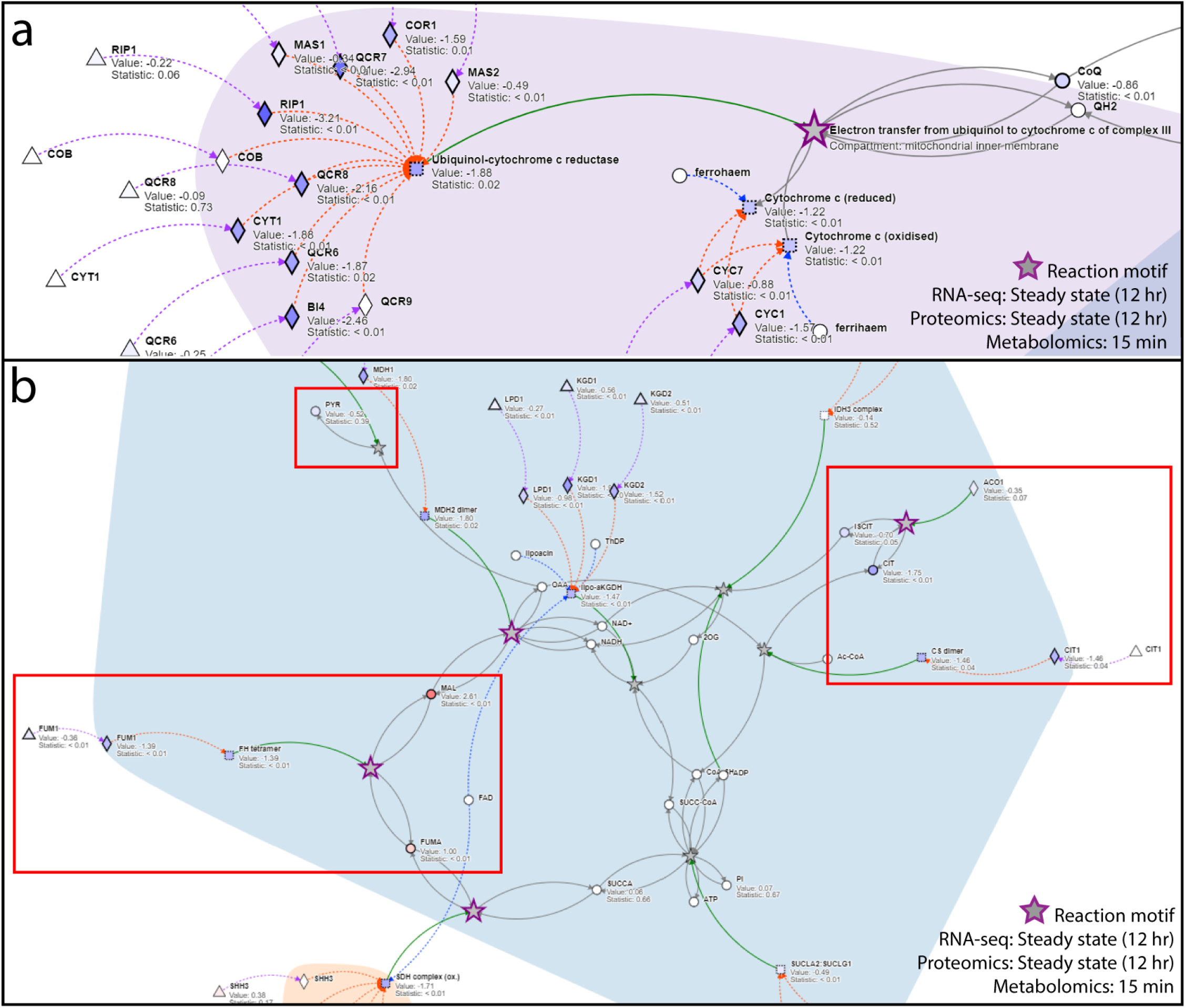
Metaboverse identifies and contextualizes important respiratory signatures in *mct1*Δ cells. **a**. Steady-state proteomics (12 hours) overlaid on the reaction, “Electron transfer from ubiquinol to cytochrome c of complex III.” **b**. Steady-state proteomics (12 hours) and metabolomics at 15 minutes overlaid on TCA-related reactions. Time stamps for each data type are displayed in the lower-right hand corner of each subplot. Measured values are shown as node shading, where an increasingly blue shade indicates downregulation, and an increasingly red shade indicates upregulation. Measured log_2_(fold change) and statistical values for each entity are displayed below the node name and represent the comparison of *mct1*Δ vs. wild-type samples. RNA-sequencing comparisons contained n=4 in each group, proteomics comparisons contained n = 3 in each group, and metabolomics comparisons contained n = 6 in each each comparison group, except for the 3-hour wild-type group, which contained n = 5. A gray node indicates a reaction. A bold gray node with a purple border indicates a potential regulatory pattern at this reaction for the given data type time points. Circles indicate metabolites, squares indicate complexes, diamonds indicate proteins, and triangles indicate gene components. Gray edges are core relationships between reaction inputs and outputs. Green edges indicate a catalyst. Dashed blue edges point from a metabolite component to the complex in which it is involved. Dashed orange edges point from a protein component to the complex in which it is involved. Dashed purple edges point from a gene component to its protein product. Protein complexes with dashed borders indicate that the values displayed on that node were inferred from the constituent protein, metabolite, and gene measurements. The background shading demonstrates Metaboverse’s ability to show cellular compartmentalization, although users may opt to toggle compartment shading off at any time.

**S9 Fig:**
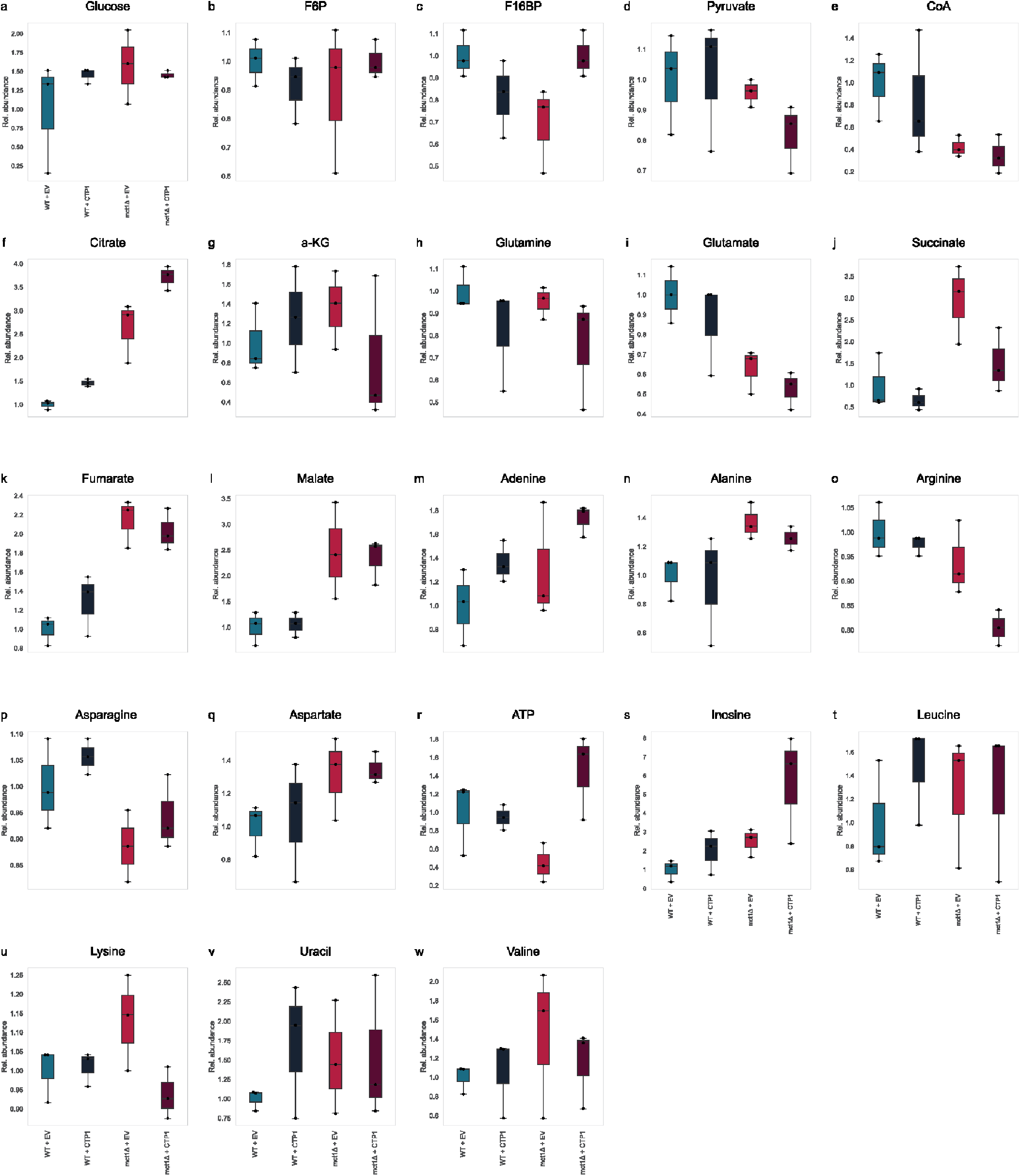
Metabolite relative abundance changes during *CTP1* overexpression in *mct1*Δ or wild-type backgrounds. Boxplot overlaid with swarm plot for metabolites quantified by LC-MS in the *mct1*Δ or wild-type background with either an empty vector or vector overexpressing *CTP1* for **a**. glucose, **b**. fructose 6-phosphate (F6P), **c**. fructose 1,6-bisphosphate (F16BP), **d**. pyruvate, **e**. CoenzymeA species (CoA), **f**. citrate, **g**. *α*-ketoglutarate (a-KG), **h**. glutamine, **i**. glutamate, **j**. succinate, **k**. fumarate, **l**. malate, **m**. adenine, **n**. alanine, **o**. arginine, **p**. asparagine, **q**. aspartate, **r**. ATP, **s**. inosine, **t**. leucine, **u**. lysine, **v**. uracil, and **w**. valine. All measurements were normalized using the average of the WT + EV samples for each metabolite. Each comparison group contained n = 3 samples. Center line represents data median, top and bottom lines represent 1.5x interquartile range. All data points are visualized as dots.

**S10 Fig.**
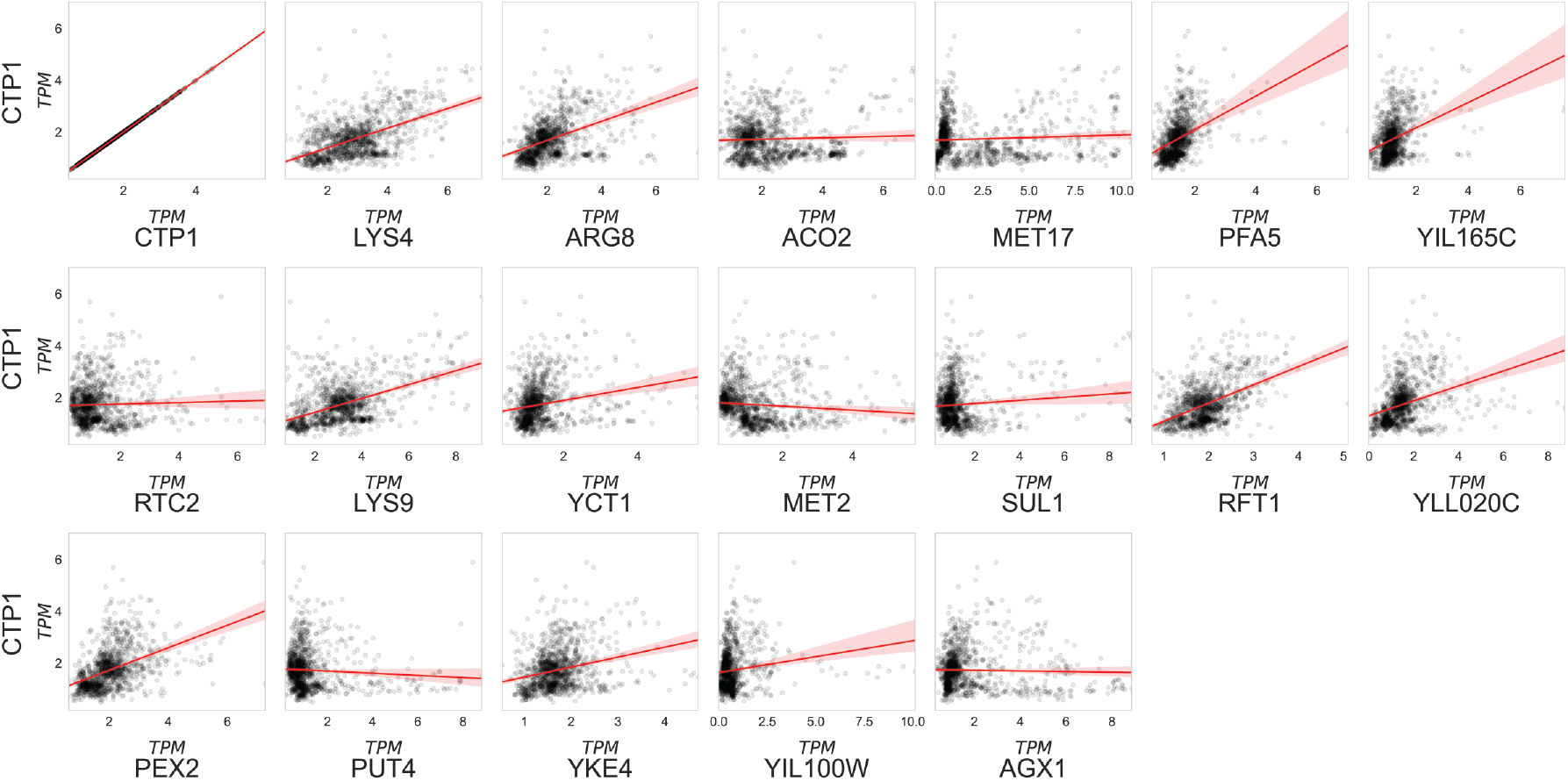
Yeast co-expression analysis of *CTP1* across all wild-type samples in refine.bio. Genes that correlated with *CTP1* (r *>* 0.5) after SpQN normalization from the wild-type samples available in the refine.bio compendium [44, 45] (n = 1248).

**S11 Fig.**
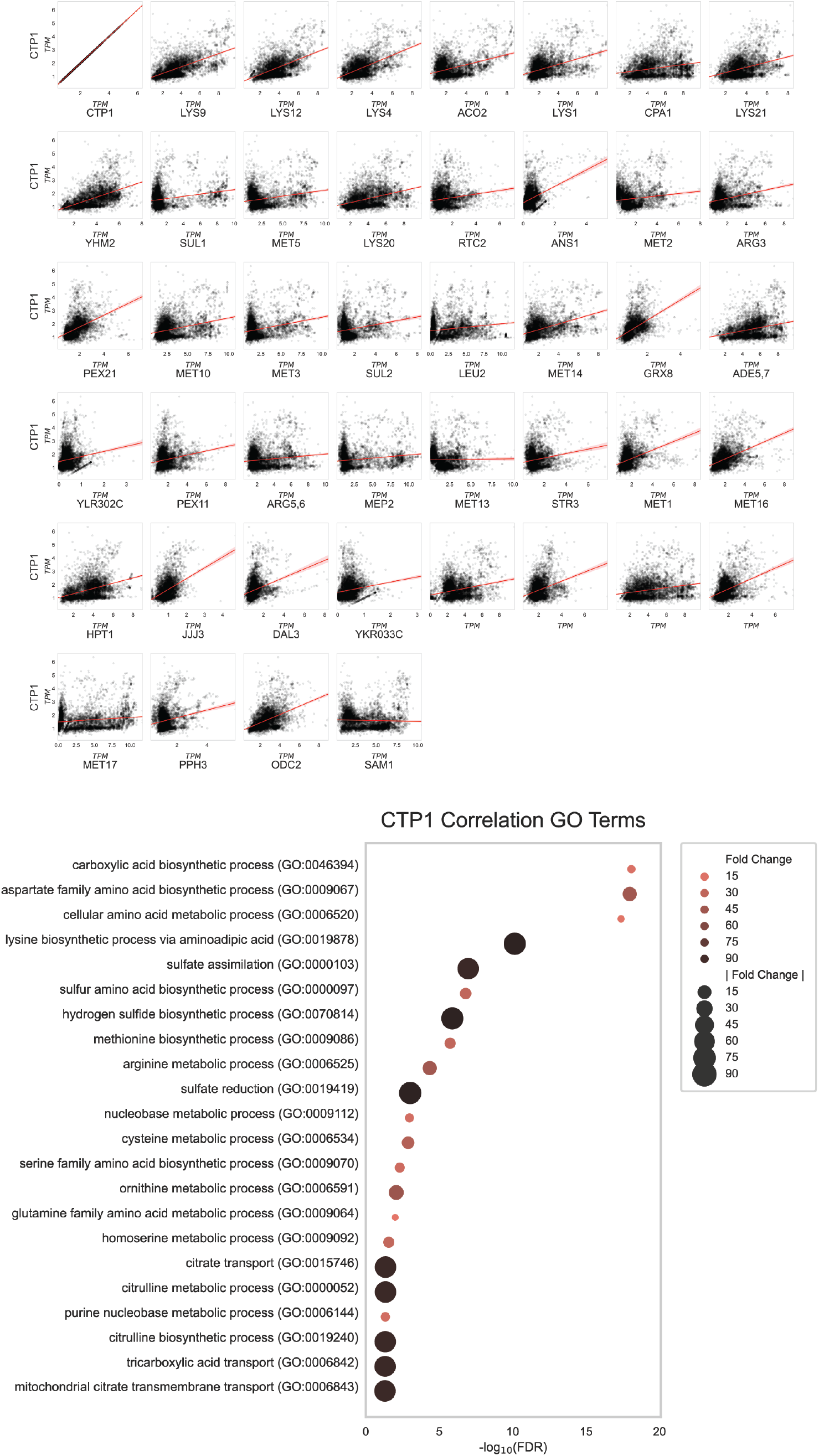
Yeast co-expression analysis of *CTP1* across all samples in refine.bio. top. Genes that correlated with *CTP1* (r *>* 0.5) after SpQN normalization from all samples available in the refine.bio compendium (n = 6370) [44, 45]. **bottom**. Bubble plot for the GO term enrichment results for genes identified in the SpQN-corrected co-expression analysis of *CTP1* across all refine.bio yeast samples. -log_10_(FDR) is plotted along the x-axis and fold change enrichment is plotted as bubble size and color intensity.

**S12 Fig:**
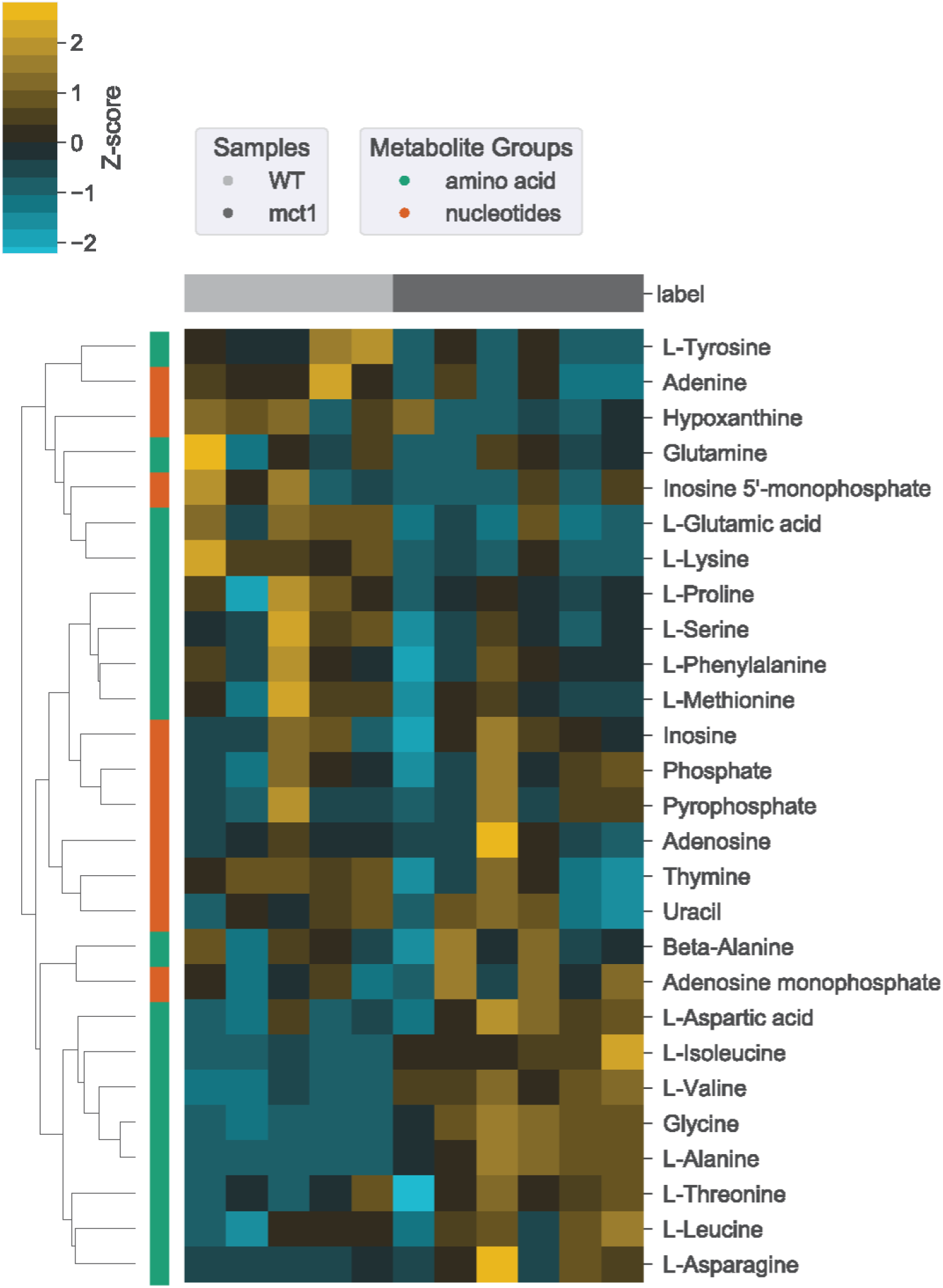
Amino acid metabolism rewiring occurs in early (3 hours) metabolomics data in the *mct1*Δ background. Heatmap of amino acid and nucleotide metabolites for wild-type and *mct1*Δ mutant strain proteomics at 180 minutes post-raffinose carbon source shift. Metabolomics comparisons contained n = 6 in each each comparison group, except for the 3-hour wild-type group, which contained n = 5. Heatmap values were mean-centered at 0 (z-score). Hierarchical clustering was performed where indicated by the linkage lines using a simple agglomerative (bottom-up) hierarchical clustering method (or UPGMA (unweighted pair group method with arithmetic mean)).

**S13 Fig:**
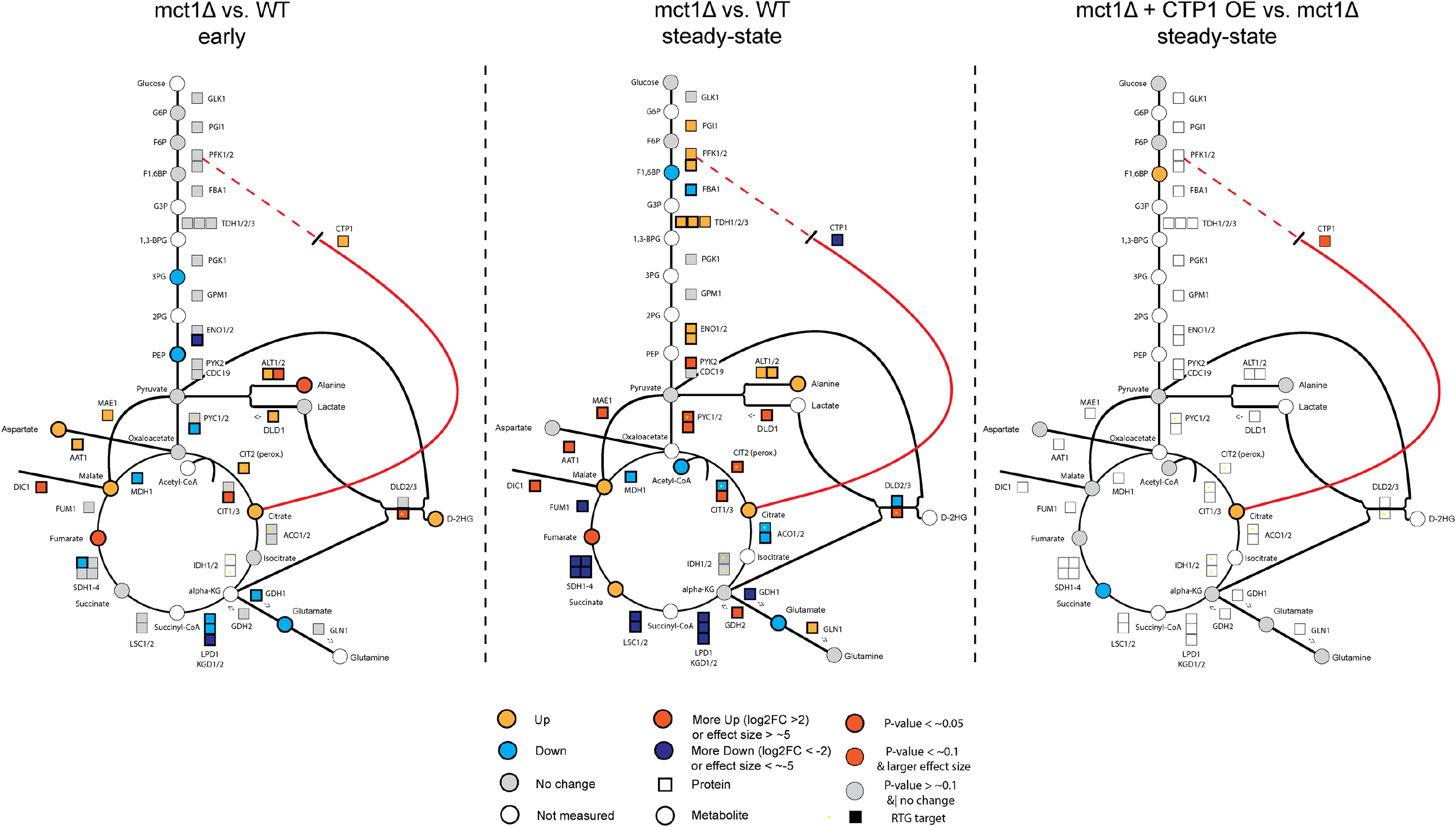
Graphical model of *mct1*Δ regulation. Graphical overview of yeast glycolysis pathway and other related reactions overlaid with summary annotations based on RNA-sequencing, proteomics, and metabolomics measurements.

**S14 Fig:**
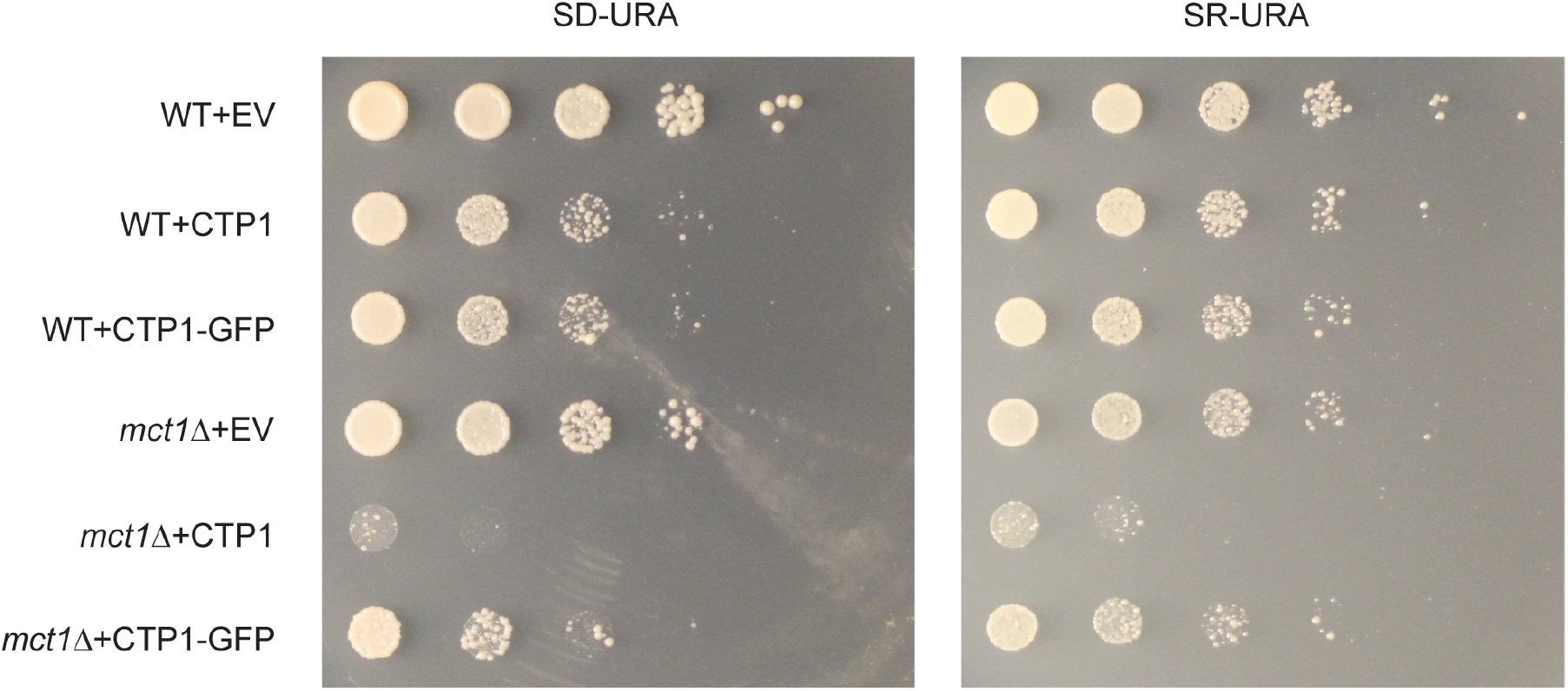
*MCT1* deletion and *CTP1* overexpression growth assays. Spot dilutions of wild-type and *mct1*Δ yeast transformed with either empty vector (EV), *CTP1* overexpression (CTP1) vector, or *CTP1*-GFP fusion overexpression (CTP1-GFP) vector on synthetic media lacking uracil supplemented with either 2% glucose (left) or 2% raffinose (right). Cells were plated at mid-log phase (OD_600_=0.3-0.6).

